# Bridging fungal resistance and plant growth through constitutive overexpression of *Thchit42* gene in *Pelargonium graveolens*

**DOI:** 10.1101/2024.03.07.583846

**Authors:** Kahkashan Khatoon, Zafar Iqbal Warsi, Akanksha Singh, Kajal Singh, Feroz Khan, Palak Singh, Rakesh Kumar Shukla, Ram Swaroop Verma, Munmun K. Singh, Sanjeet K. Verma, Zakir Husain, Gazala Parween, Pooja Singh, Shama Afroz, Laiq Ur Rahman

## Abstract

*Pelargonium graveolens* essential oil possesses significant attributes, known for perfumery and aromatherapy. However, optimal yield and propagation are predominantly hindered by biotic stress. All biotechnological approaches have yet to prove effective in addressing fungal resistance. The current study developed transgenic geranium bridging molecular mechanism of fungal resistance and plant growth by introducing cassette 35S::*Thchit42*. Furthermore, 120 independently putative transformed explants were regenerated on kanamycin fortified medium. Primarily transgenic lines were demonstrated peak pathogenicity and antifungal activity against formidable *Colletotrichum gloeosporioides* and *Fusarium oxysporum*. Additionally, phenotypic analysis revealed ∼2fold increase in leaf size and ∼2.1fold enhanced oil content. To elucidate the molecular mechanisms for genotypic cause, *De novo* transcriptional profiles were analyzed to indicate that the auxin-regulated longifolia gene is accountable for augmentation in leaf size, and ZF RICESLEEPER attributes growth upregulation. Collectively, data provides valuable insights into unravelling the mechanism of *Thchit42*-mediated crosstalk between morphological and chemical alteration in transgenic plants. This knowledge might create novel opportunities to cultivate fungal-resistant geranium throughout all seasons to fulfil demand.

## Introduction

Crops are prone to attack by many phytopathogens during their whole lifecycle, causing biotic stress. More specifically, 70–80% of plant diseases are attributable to fungi threatening worldwide agriculture, influencing crop production, and dissatisfying commercial demand (Stukenbrock and Gurr, 2023). Geranium (*Pelargonium graveolens L*′ *Her ex Aiton*) is an erect, branched herb with solid rose-scented aroma (Singh *et al.,* 2021; Lothe and Verma, 2023). The Geranium is world-known valuable crop inherited from Madagascar, Egypt, “Reunion,” South Africa, and Morocco (Rao, 2002). The secondary metabolites in Geranium Essential Oil (GEO) are commercially crucial compounds in high demand in the fragrance, pharmacology, and aromatherapy industries (Singh *et al.,* 2021). Additionally, GEO is the world’s top 20 essential oil, rich source of citronellol and geraniol, which are highly valuable alcohols found in *P. graveolens* essential oil (Lothe and Verma, 2023; Bergman *et al.,* 2020). Indeed, India produces only 20 tons/year of GEO and 661.38 US tons produced globally, whereas the national demand for geranium essential oil is ∼200 tons /year, satisfied by imports from other countries (Lothe and Verma, 2023; Mazeed *et al.,* 2023). Future market perception expects that global market demand for aromatherapy will grow by 7.7% annually from 2016 to 2026 to obtain over $8 billion in 2026 (https://www.cbi.eu/market-information/natural-ingredients-cosmetics/rose-geranium-oil).

However, various biotic stresses compromised the rose-scented geranium growth and productivity. Fungal infection caused by soil- and air-born fungal pathogens is the prime cause of yield loss, resulting in the absence of propagules after the humid and rainy season. Among the numerous fungal pathogens, *Colletotrichum* and *Fusarium* species are the primary agricultural potent pathogens contributing to crop yield losses (Kalra *et al.,* 1992). Such yield loss led to only 2% of geranium surviving in the field after the humid and rainy season. The weather favours above fungi to cause disease in well-grown *P. graveolens*. Currently, no geranium variety that is immune to fungal infection is available. Thus, alternative strategies are looked upon with high hopes. However, control measures, i.e., chemical compounds (fungicides), crop rotation, and agronomy activities, have been followed for decades to manage fungal infection (Stukenbrock and Gurr, 2023). Moreover, these approaches have limitations, whereas *Agrobacterium*-mediated genetic engineering could be a successful alternative to producing fungal-resistant *P. graveolens* plants to meet the industrial demand of GEO propagation in all seasons of north India (Luo *et al.,* 2021).

The plant produces fantastic diversity of proteins relevant to their resistance to pathogens in challenging environmental conditions (Erb and Kliebenstein, 2020; Legrand *et al.,* 1987). Among various Pathogenesis-related (PR) proteins, Chitinases are involved in plant defense mechanisms against fungal pathogens (Collinge *et al.,* 1993). Indeed, these mechanisms are not strong enough to protect the susceptible plants effectively. Therefore, mycoparasitic fungi (*Trichoderma species*) could be a rich source of genes potentially used to achieve genetically engineered superior crops against fungal pathogens (Carsolio et al., 1994). *Trichoderma spp*. Strains have been well known for their antagonistic effect on many phytopathogenic fungi such as *Rhizoctonia solani*, *Botrytis cinerea*, *Fusarium spp*., *Sclerotium* spp., (Howell, 2003).

According to the previous research article, *Trichoderma harzianum* has different chitinase genes for potent antifungal activity shown, for example, *Chit33* against *R. solani, chit42* against *B. cinerea, A. solani, R. solani,* and *Alternaria alternate* (Dana *et al.,* 2006). The *Chit42* owing its significant catalytic impact on the broad-spectrum agricultural pathogen (Jiménez-Ortega *et al.,* 2021). Moreover, several studies explained that genes encoding chitinases introduced in plant genomes elicit resistance against various fungal pathogens such as tobacco, rice, carrot, and cotton (Patil and Widholm, 1997).

By keeping the above knowledge in mind, the study aimed to develop a vector cassette having endochitinase *Thchit42* gene isolated from *T. harzianum* to introduce in *P. graveolens* via *Agrobacterium*-mediated genetic transformation to achieve resistant rose-scented geranium variety against aggressive agricultural fungal pathogen. Further, the study showed that transgenic geranium expressing constitutive endochitinase *Thchit42* gene are highly resistant to disease caused by both pathogens *Colletotrichum* and *Fusarium* species with improved yield in greenhouse trials (Fig. S1).

## Materials and Methods

### Fungal culture growth, RNA Extraction, and qRT-PCR analysis

The *Trichoderma harzianum* strain under fungal pathogen stress along with control, were cultured on potato dextrose agar supplemented with chloramphenicol. The culture incubated at 28 °C for seven days for optimal growth (Fig. S2). Further, pathogen interaction region served as a potent source of enormous chitinase expression therefore, total RNA extraction from inter-pathogen zone was carried out using an RNAeasy kit from Sigma Aldrich, and the RNA quality was evaluated using a Nanodrop spectrophotometer ND1000. High-quality RNA was scrutinized via 1% agarose gel electrophoresis, after which a concentration of 5µg per reaction was utilized for cDNA synthesis using Revert Aid Kit, Fermentas (USA) as documented by Afroz *et al*., (2024). Real-time quantitative PCR (RT-PCR) employed to analyze the differential expression of various endochitinases (*chit33, chit37, chit36,* and *chit42*) in *T. harzianum* and nucleotide sequences retrieved through GenBank database (<u>NCBI, USA) (</u>http://www.ncbi.nlm.nih.gov) (Acc. No.: X80006.1, AF525753.1, AY028421.1, and S78423.1). The qRT-PCR oligonucleotides were designed using the software primer3 tool listed in Table S2; the amplicon size was 100-150bp. In relative PCR, gene-specific primers were used at a concentration of 5 picomoles, and the optimized cDNA concentration ranged from 150-200ng/µl. Real-time PCR was performed on the 7900HT Fast Real-Time PCR system using SYBR Green Master Mix (ThermoScientific^TM^). *ThActin* was selected as the reference gene for normalizing the expression of the target gene. Relative transcript expression was determined using the 2^−ΔΔC^_T_ method (Srivastava *et al.,* 2021). Triplicate reactions were conducted for each cDNA sample to ensure data correctness.

### Molecular cloning of endochitinase *Thchit42* from *T. harzianum*

#### TA Cloning of *Thchit42* gene

The synthesized cDNA was a template for amplifying the sequence encoding *Thchit42* through semi-quantitative PCR (Fig. 1a). Each PCR reaction was 25 µl, comprising reaction buffer, 10mM dNTPs, gene-specific forward and reverse primers, and nuclease-free water added to maintain the reaction volume. PCR conditions for *Thchit42* amplification were as follows: initial denaturation at 95 ^◦^C: 5 min; 35 cycles of 95 ^◦^C: 1 min; 55.6 ^◦^C: 2 min; 72 ^◦^C: 3 min, and a final extension of 72 ^◦^C: 7 min. The amplified product underwent purification using the SureExtract® PCR Clean-up/ Gel Extraction Kit from Nucleo-pore (Genetix), and subsequently, it ligated in a pGEMT-Easy vector (Promega Madison, USA) (Fig. S3a, i). The putative recombinant white colonies were selected from a Luria agar plate fortified with IPTG as inducer, X-Gal as substrate, and ampicillin. Further, colony PCR was performed, and positive recombinant clone plasmids were extracted using the SureSpin® Plasmid Mini Kit (Genetix) (Fig. S3a, ii). To confirm the results, *Thchit42*-specific bands were obtained through restriction digestion using the enzyme XbaI and SacI (Fig. S3b, d). Sequencing was carried out with gene-specific F/R primers (Eurofins Genomics India Pvt. Ltd.) to validate the gene of interest sequence,

**Figure 1.**
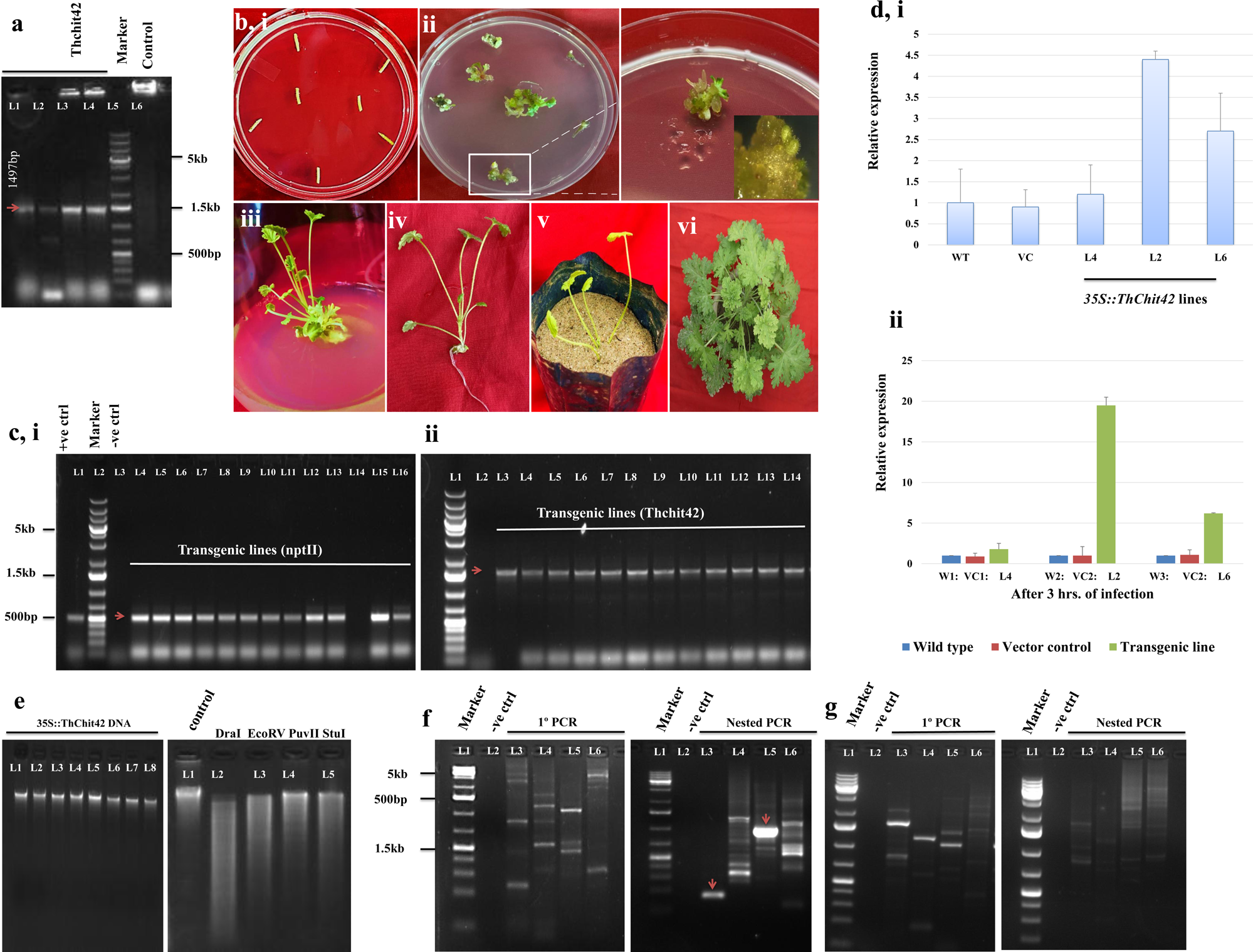
Molecular characterization of 35S::*Thchit42* in *Pelargonium graveolens*. **(a)** PCR amplicon of *Thchit42* gene (lane: L1-L4, 1497bp gene; L5-1kb + ladder; L6- −ve control. **(b, i-vii)** Regeneration and selection of *Agrobacterium*-mediated genetically transformed explants in CIM-B171. **(c, i, ii)** Twelve arbitrarily selected putative transgenic lines were evaluated through PCR with *nptII* and *Thchit42*-specific primers. +ve control (vector expression cassette); −ve control (genomic DNA of control plant); Marker-1kb + ladder. **(d, i, ii)** qRT-PCR expression analysis of putative transgenic lines indicating varying transcript levels of *Thchit42* gene pre- and post-inoculation of fungus. Experiments were performed thrice in 3 technical repeats: error bars, mean ± s.d (n=3), W-control plant, VC-vector control. **(e)** Genomic DNA of transgenic 35S::*Thchit42* line 2 and digestion with four blunt end cutters (DraI, EcoRV, PuvII, and StuI) for identification of T-DNA RB and LB flanking sequence. **(f**, **g)** Primary and nested PCR of four adaptor-ligated libraries for RB and LB flanking sequence.

### Subcloning in pBI121 vector and mobilized into *Agrobacterium tumefaciens* LBA4404 strain

Subcloning is carried out in the pBI121 vector, a commonly utilized binary vector for plant transformation. The recombinant plasmid and the empty pBI121 vector underwent digestion using the restriction enzymes XbaI and SacI (Thermo Scientific ^TM^). The purified digested insert and vector ligated using T4 DNA ligase at a 3:1 ratio. The ligation reaction mixture was then incubated at 16 °C for 18 hr., followed by enzyme deactivation at 65 °C for 30 min. Later, the transformation was initiated on an LA medium containing kanamycin as a selection marker. Milky white putative colonies were patched onto a replica plate for colony PCR using gene-specific and *nptII* primers. The plasmid from positively transformed *E. coli* (*DH5*α) was isolated and subsequently verified through restriction digestion using specific enzymes (Fig. S3c). After that, the recombinant plasmid was introduced into disarmed helper strain *A. tumefaciens* LBA4404 for plant transformation. Positive clones harboring *Thchit42* were selected on a Yeast Extract Agar plate supplemented with kanamycin and rifampicin. Recombinant clones were screened by gene-specific colony PCR, and the PCR product was analyzed on 0.8% agarose gel electrophoresis.

### *Agrobacterium*-mediated genetic transformation, selection, and phenotypic analysis of *P. graveolens* plants expressing transgenic *Thchit42*

Followed the regeneration system and standardized transformation protocol outlined by Singh *et al.,* (2021) and Horsch *et al.,* (1985). Briefly, *in-vitro* cultured petiole explants from *Pelargonium graveolens* variety CIM-BIO171 were infected with *Agrobacterium* carrying 35S::*Thchit42* for genetic transformation. After bacterial infection, the explants were inoculated onto MS medium and subjected to 24-72hr co-cultivation at 28°C in the dark (Warsi *et al.,* 2023). Subsequently, the explants were transferred to a regeneration medium consisting of 2.5 mg/L BAP, 0.1 mg/L NAA, and 1 mg/L ADS. The medium also contained 25 mg/L antibiotic kanamycin to select potential transgenic plants and 250 mg/L cefotaxime to prevent overgrowth of *A. tumefaciens*. The inoculated putative transformed explants were maintained in the culture room at 25 ± 2° C with 16h light: 8h dark photoperiod. After 40-45 days, the regenerated shoots shifted to ½ strength MS medium for root development. Subsequently, robust transgenic plants with well-developed roots turned to sterile sand for hardening.

Moreover, acclimatized plants were transferred to pots with a diameter of 10cm, filled with a mixture of vermicompost & soil in a ratio of 3:1, as shown in Fig. 1b. For phenotypic evaluation, 2-3 months putative transgenic plants acclimatized in a glass house considered. Measurements for leaf size, petiole size, inter-nodal distance, plant height, trichomes number, and biomass were recorded with three replicates for each transgenic line and the non-transgenic control (Fig. 2a, b). The conditions maintained in the glasshouse included a 12-hour photoperiod, with a temperature of 28 °C in the light and 22 °C in the dark, along with relative humidity of 70%.

**Figure 2.**
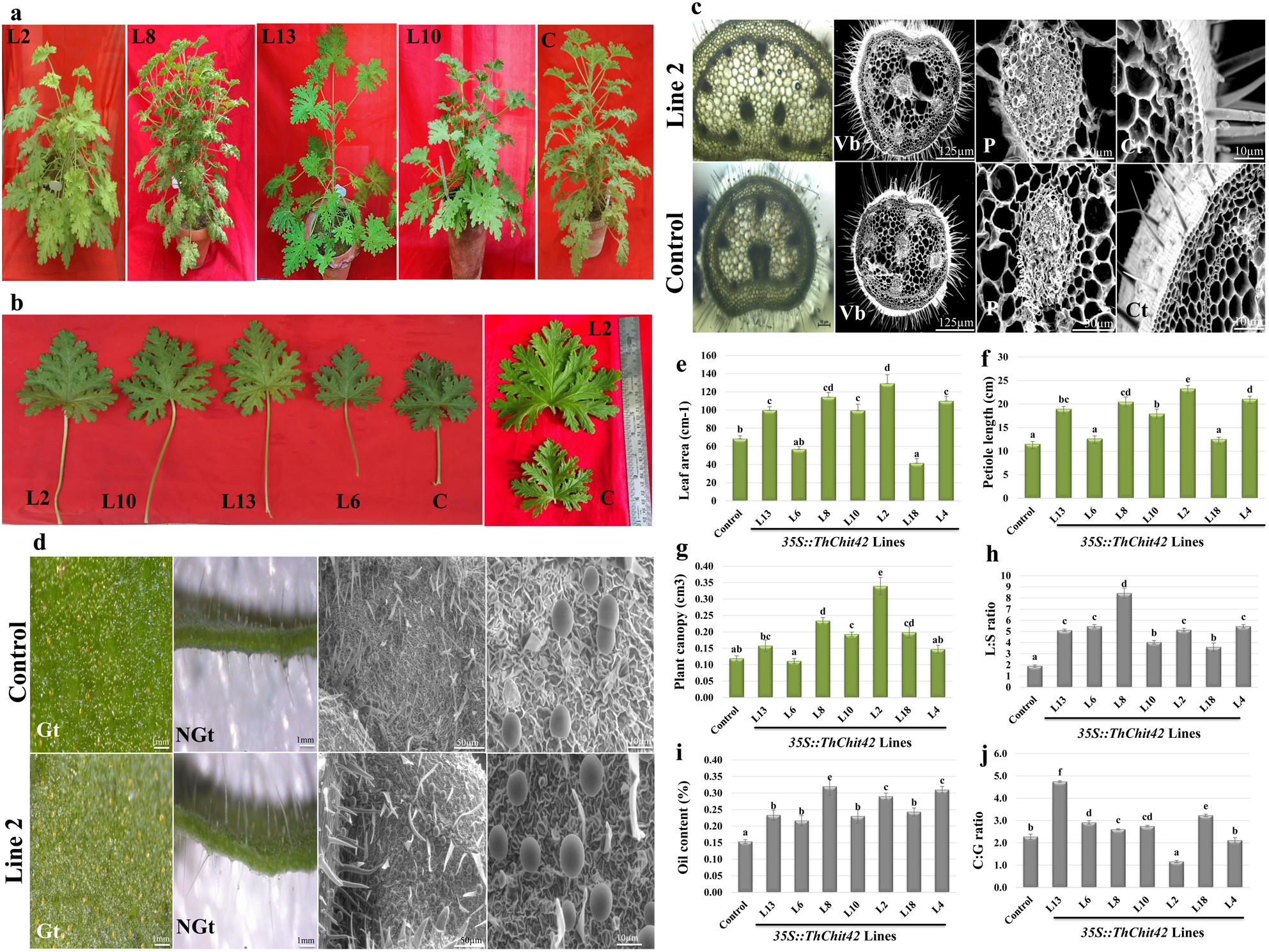
Overexpression of 35S::*Thchit42* construct and its enduring effect on phenotype and chemical composition. **(a)** Phenotypic observation between non-transgenic control (C) and different transgenic lines. **(b)** Enlarged leaf and petiole size with thickness in 35S::*Thchit42* harbouring plants compared to non-transgenic control (C). **(c)** Histological analysis reveals an elevated number of vascular bundles (Vb), increased pith (P), and cortex (Ct) size in the transgenic line (L2) in contrast to non-transgenic (C). **(d)** Amplified glandular and non-glandular trichomes (Gt and NGt, respectively) in transgenic lines and less intensification in non-transgenic plants, examined under a stereomicroscope and scanning electron microscope. **(e-g)** Size comparison of non-transgenic plants (control) and 35S::*Thchit42* transgenic lines **(e)** leaf area **(f)** petiole length **(g)** plant canopy. **(h, i)** Essential oil yield evaluation of matured transgenic lines and non-transgenic control **(h)** leaf: stem ratio **(i)** oil content (%). **(j)** Oil quality analysis based on citronellol and geraniol (C: G) of non-transgenic control and transgenic lines. Error bars, mean ± s.d. (n = 3 plants).

### Molecular analysis of potential transgenic lines expressing *Thchit42*

#### PCR and Differential gene expression through Real-Time quantitative PCR (RT-PCR)

Twelve arbitrarily selected putative transgenic rose-scented geranium lines were evaluated for transgene *Thchit42* stable integration into genomic DNA for molecular analysis. The genomic DNA was isolated using the CTAB method. The quality of genomic DNA was assessed using Nanodrop spectrophotometer ND1000 in preparation for PCR analysis. The gene-specific PCR reaction is described in the cloning section of online methods. To verify the occurrence of construct integration through homologous recombination, selection marker-specific amplification was conducted following the protocol outlined by Warsi *et al*., (2023). *Thchit42* and *nptII* amplified products observed in 0.8% gel electrophoresis containing 5µl/100ml EtBr. The relative gene expression of *Thchit42* positive lines was confirmed through RT-PCR. Oligonucleotides used for RT-PCR analysis were designed using Primer3, an online available tool and listed in Table S2. Total RNA was extracted using spectrum ^TM^ Plant total RNA kit (Sigma Aldrich), and purified RNA was further preceded for cDNA preparation using Revert Aid First strand cDNA synthesis kit (ThermoScientific ^TM^). Quantitative RT-PCR was conducted in 7900HT Fast Real-Time PCR using SYBR Green Master Mix from Thermo Scientific ^TM^. The reference gene used to normalize target expression data was *PgActin*, and transcript expression was analyzed by following the 2^−ΔΔC^_T_ method (Fig).

### Genomic DNA Walking

To identify an unknown sequence adjacent to the integration site of the T-DNA construct, a genomic DNA walk was carried out using the Universal Genome Walker 2.0 Kit (Takara Bio Inc. Company, USA). Genomic DNA was isolated using DNeasy ® Plant Mini Kit (Qiagen), and quantity was measured using a Nanodrop spectrophotometer ND1000. The optimum DNA quantity to prepare genome walking libraries employs four restriction enzymes: DraI, EcoRV, PvuII, and StuI. Subsequently, each purified individual library is ligated to the genome walker adaptor. The ligated product of each library was employed for both primary and nested PCR, followed by the visualization of the resulting product on a 1.5% gel electrophoresis to achieve PCR-based DNA walking (Fig. 1f, g). The adaptor primers 1 and 2 are provided in the kit. At the same time, 5’ Pnos and 3’ Tnos oligonucleotides were prepared for primary and nested PCR using the OligoAnalyzer and PCR primer stat tool, as shown in Table S3; Fig. S5. The secondary PCR product was cloned in pGEMT-Easy vector (Promega Madison, USA), and Sequencing was performed using the nos promoter’s AP2 and 5’ junction (Eurofins Genomics India Pvt. Ltd.).

### Microscopic evaluation of heterologous expressing transgenic lines

Two months of transgenic and non-transgenic control plant samples were used for histological and histochemical analysis. Samples were prepared for imaging under Leica Microsystem limited version 2.1.0 (Switzerland), fluorescence microscopy, and Environmental scanning electron microscope (ESEM) to analyze histological information of cell wall thickening, number of vascular bundle, lignification, ROS production, and trichome density (Fig. 2c,d).

### Essential oil extraction and chemical composition of transgenic *Thchit42* lines

3-4 months old, seven healthy acclimatized glass house transgenic lines and WT used for oil analysis. As per the Majeed *et al*., (2023) reports, ∼ 100g of fresh sample was weighed and subjected to hydro-distillation with 500 ml of D.W. using a Clevenger-type apparatus for 180 min. The oil yield is calculated using the formula oil yield = (essential oil * fresh weight)/100. Furthermore, for the identification of GEO composition, GC-MS was performed (Upadhyay et al., 2022).

### Isolation procedures, pathogenicity bioassays, and assessment of antifungal activity

#### Isolation of fungal pathogens

To investigate the pathogenicity and antifungal activity, fungal pathogens responsible for significant infection in *P. graveolens* were isolated from diseased geranium plants (CSIR-CIMAP, Lucknow). Initially, diseased tissue samples were rinsed with sterile double-distilled water (SDW), followed by treatment with a disinfectant solution containing 75% ethanol and 1% sodium hypochlorite, each for 30 seconds. Subsequently, the samples were triple-washed with SDW. Treated tissue samples (1×1 cm2) were then cultured on Potato Dextrose Agar (PDA) plates and incubated for seven days at 28℃ to facilitate the isolation of associated fungi as per the methodology outlined by Jogee *et al*., (2017). Furthermore, *Fusarium oxysporum* and *Colletotrichum gloeosporioides* isolated from root and leaf samples, respectively, were purified. Following purification, fungal spores were observed using both a light microscope (LeicaDM750, Germany) and Scanning Electron Microscopy (SEM) (FEI Quanta 250 SEM Netherland) (Figure 3b, c). Koch’s postulate followed to validate the isolated fungi’s pathogenicity.

**Figure 3.**
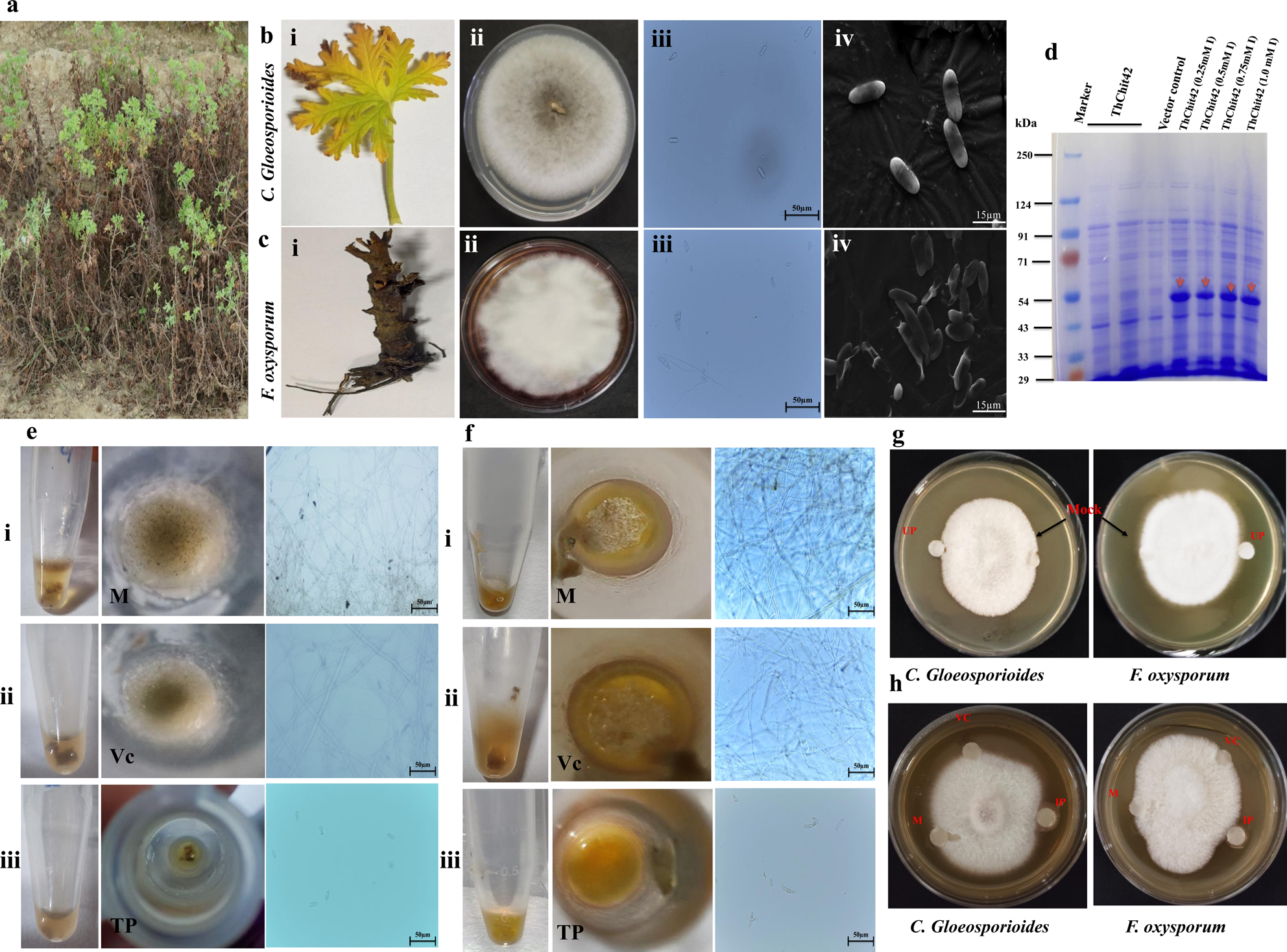
Isolation of potent fungal pathogen and antifungal activity. **(a)** Infected *Pelargonium graveolens* (non-transgenic) during the rainy and humid season. **(b, i)** Isolation of vigorous *Colletotrichum gloeosporioides* from diseased *P. graveolens* leaf **(b, ii-iv)** Morphological appearance of *C. gloeosporioides* and magnified view of spores, examined under (**iii)** Light microscopy, scale bar-50 μm; **(iv)** Scanning Electron Microscopy (SEM), scale bar-30 μm. **(c, i)** *Fusarium oxysporum* extraction from Infected stem. **(c, ii-iv)** Morphological analysis maximizes the view of spores, viewed under **(iii)** Light microscopy, scale bar-50 μm; **(iv)** Scanning Electron Microscopy (SEM), scale bar-30 μm. **(d)** SDS-PAGE analysis of *ThChit42* protein expression under un-induced and IPTG-induced conditions. **(e, f)** Antifungal activity test against *C. gloeosporioides* and *F. oxysporum,* respectively. **(e, f, i)** Mock (M) -aggressive mycelial growth. **(e, f, ii)** Vector control (Vc) total protein - optimum mycelial growth. **(e, f, iii)** recombinant total protein (TP)-significant mycelial growth inhibition; all mycelial growth patterns examined under the light microscope, scale bar-50 μm. **(g, h)** Antifungal activity by disc diffusion method against *C. gloeosporioides* and *F. oxysporum*; M-mock; IP-induced protein; VC-vector control; and UP-un-induced protein. Experiments were conducted in triplicates with three technical replicates.

### Plant infection and pathogenicity bioassays

Both fungal isolates were cultured on PDA supplemented with chloramphenicol at 28^D^C for seven days within a BOD incubator to investigate disease resistance levels. Subsequently, 13 lines expressing 35S::*Thchit42* were screened for antifungal bioassays (Table S4), for both *in-vitro* leaf detached and *in-vivo* bioassay, leaves and roots punctured to inject conidial suspensions (10^8^ conidia/ml) via a sterile needle that notably evaluated the pathogenicity of *C. gloeosporioides* and *F. oxysporum,* respectively. As a control, SDW was injected in place of spore suspension. Plants enveloped in transparent plastic sheets to maintain high relative humidity (RH) were placed in a growth chamber at 28^D^C with a 12-hour photoperiod for six days after pathogen injection (DPI) to evaluate the infection rate. Subsequently, the disease severity (DS) index method was employed to rank transform plants alongside the non-transgenic control after 7 DPI (Guarnaccia *et al.,* 2019). The pathogenicity severity was categorized on a scale of 0 for healthy plants to 100 for severe instances featuring necrotic patches and deceased plants. Specifically, a score of 25 indicates mild virulent lesions with slight leaf chlorosis, 50 represent the moderate presence of characteristic symptoms, and 100 represents a severe manifestation. The pathogen aggressiveness for *P. graveolens* was categorized as follows: Low L (disease index ≤25); Average = M (disease index 26–50); and High H (disease index > 51). The inoculated fungi were re-isolated and identified by Koch’s postulates to confirm the result.

### Histochemical quantitative analysis of Reactive Oxygen Species (ROS) production

Intensity of H_2_O_2_ and O_2_^−^ has evaluated through 3, 3’-Diaminobenzidine (DAB) and Nitro Blue Tetrazolium (NBT) respectively as per Warsi *et al*., (2023) on fungal stress non-transgenic control and transformed plants (Fig. S9).

### *Thchit42* expression and antifungal activity

To assess the antifungal activity, the mycelia of *F. oxysporum* and *C. gloeosporioides* were cultivated on a PDA plate to evaluate total protein activity. Subsequently, *Thchit42* was cloned in expression vector pET28a (+) to generate a construct for 6x His-tagged recombinant protein expression. This engineered protein was induced in *E. coli* using 0.25mM IPTG and maintained at 16°C overnight. The antifungal assay used total protein extract derived from the bacterial expression system. Two different approaches were implemented for the evaluation of antifungal activity. In the first approach, *F. oxysporum* and *C. gloeosporioides* spores (10^8^ conidia/ml) inoculated in PDB amended with total protein isolated from the recombinant expression bacterial system. MilliQ water was used in the control. After five days of incubation at 28°C, mycelia growth and spores in each set were recorded using light microscopic (LeicaDM750, Germany) (Fig. 3e, f). In the second approach, sterile filter paper discs (0.6 mm diameter) containing total protein served as the experimental treatment, empty vector as the control, and Mock. The prepared filter paper discs, representing the respective treatments, were placed on either side of the fungus centrally placed within a 90 mm petri plate. The sterile filter paper disc moistened with deionized water served as a Mock (Schlumbaum *et al*., 1986). This experiment configuration allows for examining the localized effect of the total protein extract on fungal growth and spore formation. Using sterile filter paper discs facilitates the controlled delivery of experimental and control treatments of the impact on the target organism. Further, image analysis of the antifungal assay replications was conducted using Adobe Photoshop 7.0. The objective was to analyze the average inhibition area of mycelia growth for both pathogenic fungi.

### *De novo* transcriptome analysis

For De novo transcriptome analysis, three leaf samples of the same age and weight were pooled together for RNA isolation (for each sample, transgenic and non-transgenic control). Further, the transcript obtained from mRNA processed for quality check then the raw reads pre-processed using FastQC v.0.11.9 toolkit and Cutadapt v.3.4. The high-quality reads assembled *de novo* by using Trinity v2.14.0 assembly software and assessed by N50 statistics, reads-remapping percentage (Bowtie2), and mapping against the orthologous database using BUSCO v.5.4.5 (Benchmarking Universal Single-Copy Orthologs). BlastX for the homology search of the non-redundant transcriptome obtained using CD-HIT-EST v4.6.1 against NCBI Non-redundant (Nr) database with E-value cut-off of 10^−5^. The G.O. terms assigned to the blasted sequences utilized for mapping biological pathways against the Kyoto Encyclopedia of Genes and Genomes (KEGG). Transcripts were quantified using RSEM (RNA-seq Expectation Maximisation), and cross-sample normalization was done using TPM (transcript per million). Differentially expressed transcripts (DEGs) were identified using the edgeR package with log fold change (logFC) ≤−2 for downregulated or ≥2 for upregulated, respectively, and FDR (false discovery rate) of <0.05. Pathway & G.O. enrichment analysis was performed using the Fisher exact test with Benjamini–Hochberg false-discovery rate (FDR) < 0.05.

### Statistical Analysis

The analysis of recorded data was done using SPSS v17.0 statistical software. Data were subjected to ANOVA, and means were compared using the least significant difference (LSD) test at a 0.05 probability level. The ANOVA was used to evaluate plant phenotypic and chemical composition of transformed *Pelargonium graveolens* lines along with control on the recorded parameters, using a completely randomized design (CRD) for an overall effect of *Thchit42*. For the comparison of means, a post hoc test was carried out using Tukey HSD with the help of SPSS statistics 17.0 at a 95% confidence level (P < 0.05).

Plant canopy = 2/3 pi H (A/2 x B/2)

Where H = plant height, A = A canopy diameter (m) in E-W, and B-canopy diameter (m) in N-S direction.

### Research materials

*Trichoderma harzianum*, *Fusarium oxysporum,* and *Colletotrichum gloeosporioides* fungal Strains obtained from CSIR-CIMAP for lead gene *Thchit42* selection and cassette development. *Pelargonium graveolens* (CIM-BIO171) was taken from CSIR-CIMAP field to produce fungal resistant transgenic plants. *Thchit42* expression cassette and fungal resistant transgenic lines are available from CSIR-CIMAP.

## Result

### Construct formation and genetic engineering

#### Mining of the target gene and molecular cloning of *T. harzianum* endochitinase *Thchit42*

Quantitative RT-PCR performed to identify the target gene among all the illustrious endochitinases of *T. harzianum,* indicating notably higher relative transcript expression for *Thchit42* under pathogen stress, as per the 2^−ΔΔC^_T_ value (Fig. S2). Therefore, the entire coding sequence of *Thchit42* is amplified using fungus complementary DNA (cDNA) as a template. Subsequently, the amplified PCR product was inserted into the binary vector pBI121, between CaMV35S and nopaline synthase terminator, as shown in Fig. S3a-d. The amplified PCR product was also cloned in pET28(+) plasmid for expression of 6x His-tagged recombinant protein in *E. coli* (Fig. 3d).

### Regeneration, selection, and molecular evaluation of putative genetic transformants

Genetic transformation initiation involved the integration of the target gene into the T-DNA region of *Agrobacterium tumefaciens* strain LBA4404 through homologous recombination. ∼120 Putative transgenic shoots selected on a kanamycin-supplemented regeneration medium. Regenerated chimeric shoots removed through five rounds of subculturing on the selection medium (Fig. 1b, i-iv). Healthy, rooted, putative transformed rose-scented geranium lines were transferred to glass house for acclimatization (Fig. 1b, v-vi). Remarkably, the survival rate during the hardening procedure for putative transgenic plants was 70.0 ± 0.9%. Molecular characterization of 12 healthy transgenic *P. graveolens* lines was conducted to ascertain the presence of selection marker (*nptII)* and gene-specificity (*Thchit42*) (Fig. 1c). To validate these results, qRT-PCR was carried out both pre- and post-infection with the fungus *C. gloeosporioides.* Notably, the relative transcript expression of the *Thchit42* gene exhibited a substantial increase, ∼ 5-fold higher after 3 hours of infection compared to the non-infected state (Fig. 1d). Collectively, these results confirmed the successful integration of the *Thchit42* gene into the plant genome, with the upregulation of “*Thchitinase*’ under fungal stress conditions.

### Pathogenicity bioassays and antifungal activity assessment for disease resistance

Pathogenicity bioassays were performed to predict the disease severity index of transformed 35S::*Thchit42* lines compared to non-transformed control. Control and transformed plants infected by applying spore suspensions of *C. gloeosporioides* on leaves and *F. oxysporum* on root tissues. Subsequent evaluation revealed that only subset of transgenic lines indicated a disease severity index of 0, scoring at seven days post-inoculation (DPI), with line L2 demonstrating the most robust fungal resistance response (Fig. 4). Conversely, the WT *P. graveolens* showed characteristic circular or irregularly shaped lesions on leaves that merged to form larger affected areas in the case of *C. gloeosporioides* (Fig. 4a, i). Similarly, roots infected with *F. oxysporum* exhibit discoloration, typically appearing brown or reddish-brown, compared to healthy transgenic line roots (Fig. 4b, i). Notably, the observed symptoms closely mirrored those encountered in the field from which the diseased plant was sourced (Fig. 3a). Koch’s postulates were rigorously fulfilled to confirm the identity of the re-isolated fungi. No symptoms were observed in the uninjured and milli Q water-injected control plants. An antifungal activity assay was conducted to substantiate the results further, revealing that total protein exhibited significant inhibition of both C. *gloeosporioides* and *F. oxysporum* mycelial growth. The degree of mycelial growth inhibition for both fungi positively correlated with increasing total protein (Fig. 3h). Additionally, the inhibitory impact of the complete protein is more pronounced for *F. oxysporum* than *C. gloeosporioides* (Fig. 3g, h). The results were further validated by the PDB broth assay, wherein inhibition of mycelia and spore germination was observed in the control set (Fig. 3e, f). The result concludes, line L2 indicated a significant disease severity index and expression of *Thchit42* has antifungal activity against both *F. oxysporum* and *C. gloeosporioides*.

**Figure 4.**
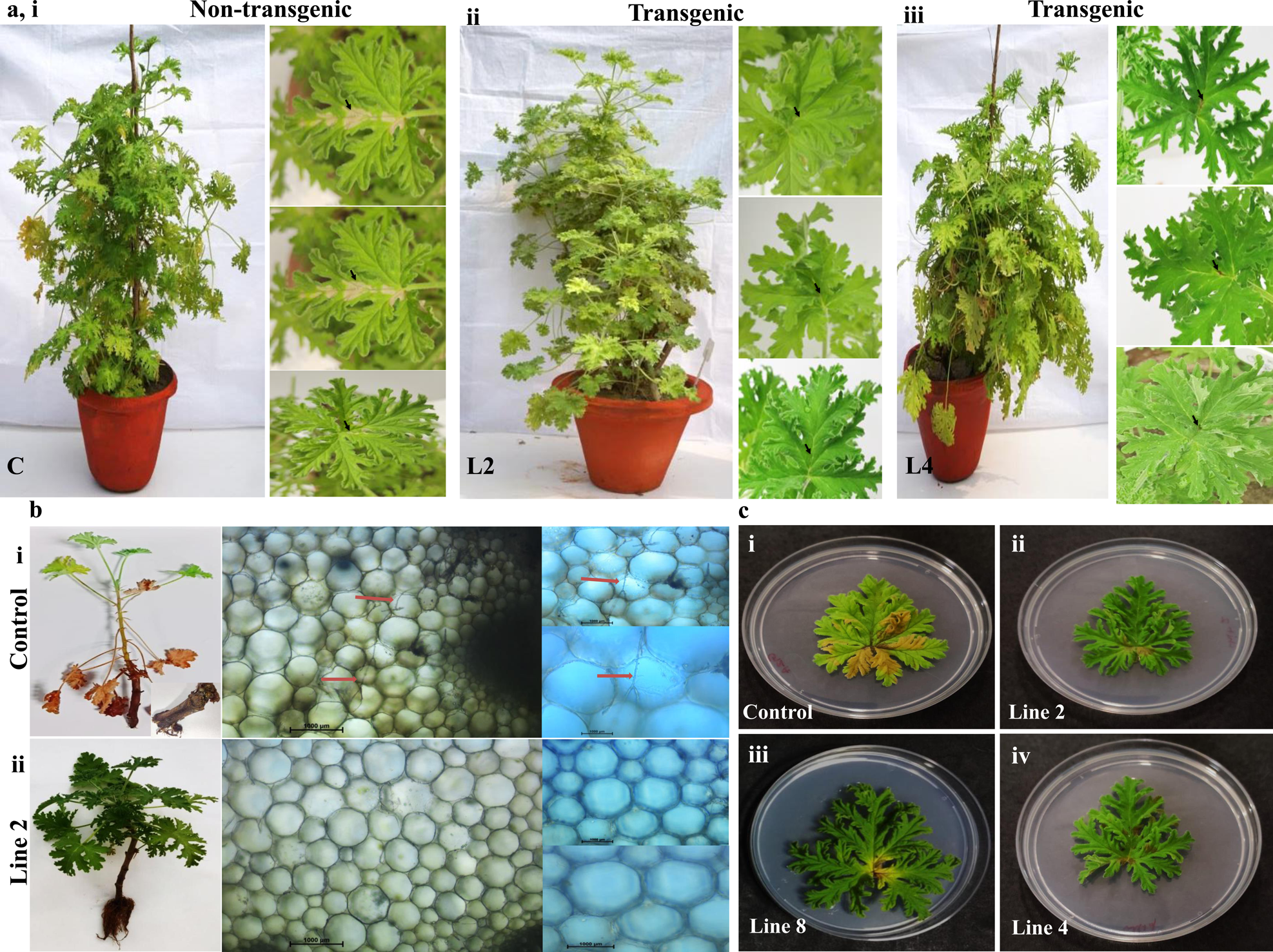
Pathogenicity bioassay evaluations of transgenic and non-transgenic plants for disease resistance scoring. **(a, i-iii)** *In vivo,* pathogenicity bioassay of *C. gloeosporioides* showing different degrees of infection on **(i)** non-transgenic control (C) and **(ii, iii)** transgenic lines L2 and L4 *P. graveolens* leaves after seven days of inoculation. **(b, i, ii)** *In vivo* screening of diseased score caused by *F. oxysporum* in **(b, i)** *P. graveolens* non*-*transformed control (control) in contrast to **(b, ii)** transformed lines L2; examined under a stereomicroscope. The arrow denotes the presence of fungal mycelia in the control root tissue, while no such observation was recorded in the root section of the L2 line. **(c)** *In vitro* leaf detached assay, screening of *P. graveolens* detached leaves after seven days of inoculation, **(c, i)** non-transformed control with **(c, ii-iv)** transformed lines L2, L8, and L4 against *C. gloeosporioides*. Experiments were performed in 3 batches, each triplicating with three technical replicates.

### The Stable overexpression of the *Thchit42* heterologous gene and its enduring influence on both the phenotype and chemical composition

For phenotypic and chemical composition screening of *Thchit42* stable expression, seven healthy transformed plants were phenotypically screened for morphological distinction. Notably, the phyllome and petiole dimension in the transformed lines was significantly increased compared to the non-transformed control plant (Fig. 2a, b, e-f). Additionally, Stereo and Scanning Electron microscopic of the leaf surface revealed a conspicuous augmentation of trichomes in the transformed geranium lines in comparison to the non-transformed control plant (Fig. 2d). The leaf and petiole tissues of the transformed lines showed velvet-thickened texture on the surface. A histological study revealed an increment in no. of vascular bundle, which found an apparent correlation with increase in petiole girth (Fig. 2c). Analysis of aromatic essential oil composition and content (%) revealed, oil content enhanced to 1.4-2.1 fold. Furthermore, the fresh weight data of geranium depicted that leaf-to-stem ratio (L:S ratio) elevated ∼1.9-4.4 fold, as illustrated in Fig. 2g-h, i. The observed increase in oil content is quite reasonable as leaf size has increased after integrating the *Thchit42* gene. In addition, the GC-MS analysis revealed C: G ratio varied from 1.1-5.48% across different *P. graveolens* transformed lines (Fig. 2j; Fig. S4). In summary, the finding strongly indicates a correlation between the expression of *Thchit42* and the growth patterns and the activation of the phenylpropanoid pathway in the examined geranium plants.

### Localization of *Thchit42* gene construct in the transgenic line

A thorough investigation was undertaken through genome walking, aimed at identifying the RB and LB flanking T-DNA sequence to investigate the precise integration site of edited T-DNA (Fig. S5a). Four different libraries constructed using four restriction enzymes which were further utilized for PCR-based genome walking (Fig. 1e). Later on, in the mentioned procedure (Clonetech, Takara Bio Inc., USA), the secondary PCR products with the sizes of 0.1 and 0.9 kb successfully obtained from libraries digested through DraI and PvuII respectively from RB (Fig. 1f) and no specific bands obtained from LB flanking region (Fig. 1g). Following this, the eluted products were subjected to cloning and sequencing using AP2 and Pnos2 oligonucleotides. Moreover, performing Blastn analysis, the derived sequences exhibited no significant alignment with extended coding regions in any of the six frames. Therefore, the result concludes that the T-DNA construct has integrated at the non-coding region, as it has multiple stretches of T and A along with the TATA box (Note S1). The result strongly supports that the T-DNA integration event did not induce insertional mutagenesis, as no disruption of the coding region was identified in the analyzed sequences.

### Effect of *Thchit42* on *P. graveolens* genotype and its validation

In the context of forward genetic investigation, large-scale RNA sequence data of 35S::*Thchit42* geranium revealed genes responsible for phenotypic variation in transformed *P. graveolens* line L2 (Fig. 6; Table S1). The *Thchit42* gene, PR protein, prompts the hypothesis that its constitutive overexpression leads to the activation of the plant defense mechanism. Comprehensive profiling was performed to understand the involvement of protein and its crosstalk (Fig. 6; Fig. S8). Subsequently, the heatmap of DEGs assessment revealed the over-expression of other PR proteins (*Pgchitinase, PgEndochitinase* - yet not reported in *P. graveolens*), peroxidase, and disease resistant Response (*PgDirigent*) genes (Fig. 5a, b, d-f). In addition, DEGs data revealed upregulation of Shikimic acid and phenylpropanoid pathway (Fig. S6), which resulted in a significant increase in indole-3-acetic acid (*Pg*IAA) and flavin-3-monooxygenase (*PgYUC*) (Fig. 5h, n). Moreover, Auxin transport and signaling overexpressed, synergistically enhancing the auxin-regulated *’*Pglongifolia protein’ which strongly suggests an accountable gene for the increase in leaf size (Fig. 5g). Likewise, the expression level of PAL, coumaryl synthase, and caffeic acid-3-o-methyltransferase detectably rose (Fig. 5a; Fig. S7), leading to the cell wall thickening and lignification, (secondary defense mechanism) observed in the leaf and petiole of transformed plants. On the other hand, the upregulation of Fatty acid desaturase, Lipoxygenase, and Allene oxide synthase (*PgAOS*) was observed to increase Jasmonic acid (JA) content, boosted through up-regulation of TIFY for downstream signaling transduction, suggesting a concerted effort to enhance trichome size and number (Fig. 5a, l, q). Furthermore, MYC and WRKY TFs were upregulated notably to participate in the defense mechanism (Fig. 5a, j). Interestingly, on the other hand, primary metabolism, i.e., nitrogen metabolism, TCA cycle, carbon fixation, and Zinc finger RICESLEEPER enhancement, revealed strong correlation with plant growth and development (Fig. 5a, k, i). Expressly, glutamate synthase intensive expression has confirmed crosstalk with ethylene signaling by ERF/AP2 regulation (Fig. 5a, m, p). Among the phenotypic changes, the elongation of the petiole was due to lower expression of AT-hook motif nuclear-localized TF (AHL) and COP1, repressor of PIF-regulated genes (Fig. 5a, o). These results are considered for further validation through qRT-PCR. Therefore, the above result suggests that the *Thchit42* gene potentially co-relates fungal stress resistance and plant growth (Fig. S8).

**Figure 5.**
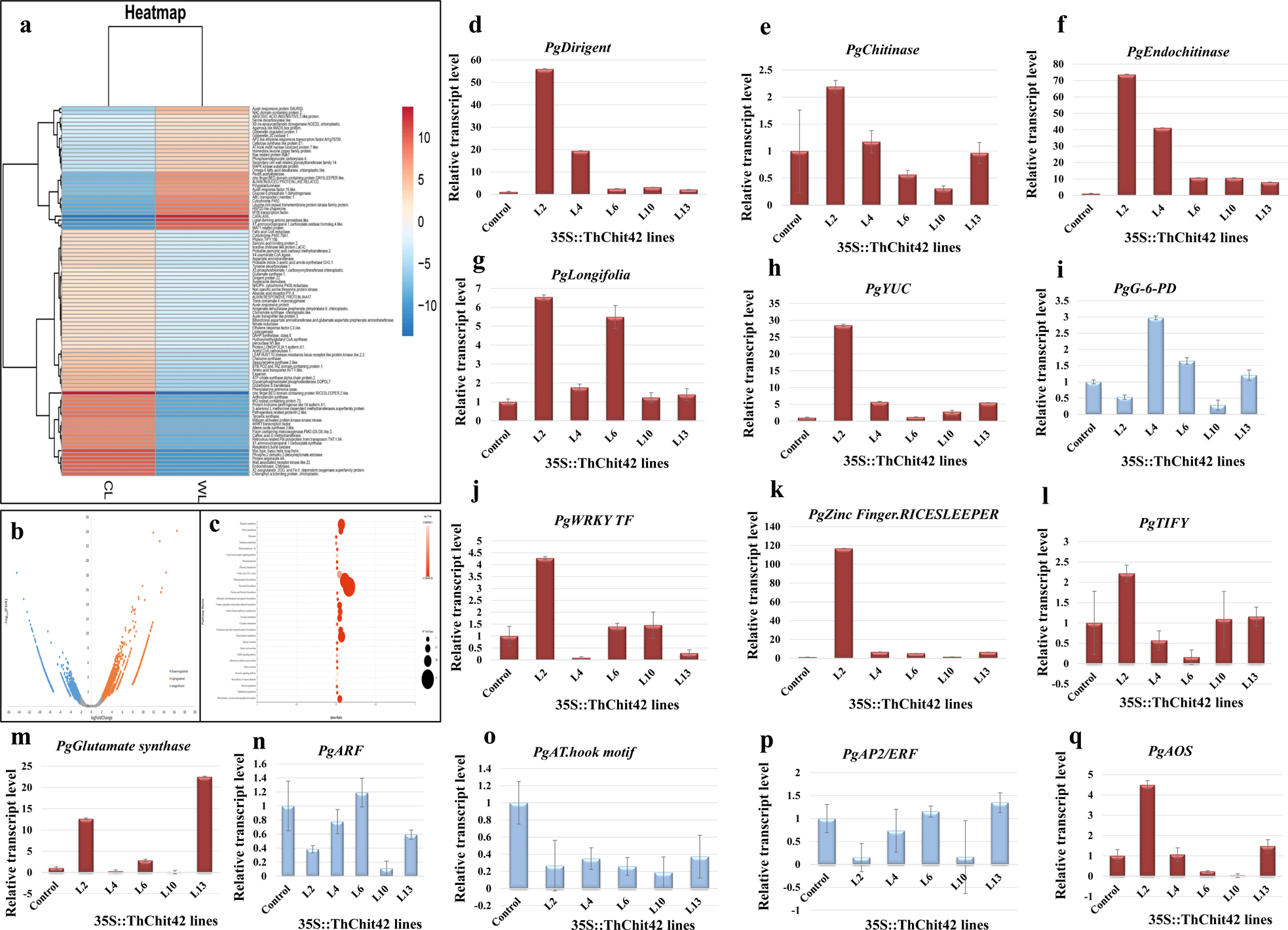
Effect of *Thchit42* on *P. graveolens* genotype and its validation. **(a**) Expression heatmap of DEGs regulating defense, growth, and phenylpropanoid pathways; WL-non-transgenic control; CL-transgenic line L2. **(b)** Volcano plot for DEGs with logFC on the x-axis and –log10 FDR on the y-axis with logFC > 2 for upregulated and logFC < −2 for down-regulated and −2 < logFC < 2 for insignificant transcripts. **(c)** Significantly enriched pathways based on FDR < 0.05. **(d)** mRNA expression level analysis of *PgDirigent* gene (disease resistance response) between non-transgenic control and transgenic lines. **(e, f)** Enhanced relative transcript expression fold of *Pgchitinase and Pgendochitinase.* **(g, h, n)** Auxin regulated longifolia, YUC, and ARF transcript expression fold change. **(i)** Differential gene expression of glucose-6-phosphate dehydrogenase (G-6-PD) through qRT-PCR. **(j, l, q)** Real-time PCR data revealed a defense mechanism promoted through enhanced log fold change of *PgWRKY, PgTIFY,* and *PgAOS.* **(k)** Enormous transcript level of Zinc finger BED domain. RICESLEEPER transposons validate plant growth and development. **(m)** qRT-PCR analysis of glutamate synthase impact on N_2_ metabolism to regulate growth. **(o)** Down mRNA expression of AT-hook motif nuclear-localized transcription factor regulates PIF expression. **(p)** AP2/ERF down-regulated relative expression level. qRT-PCR experiments performed in triplicates with three technical replicates: Error bars – mean ± s.d. (n = 3); *PgActin* considered as a reference gene for normalization; non-transgenic control mRNA considered as control and five 35S::*Thchit42* transgenic lines transcript expression was analyzed.

**Figure 6.**
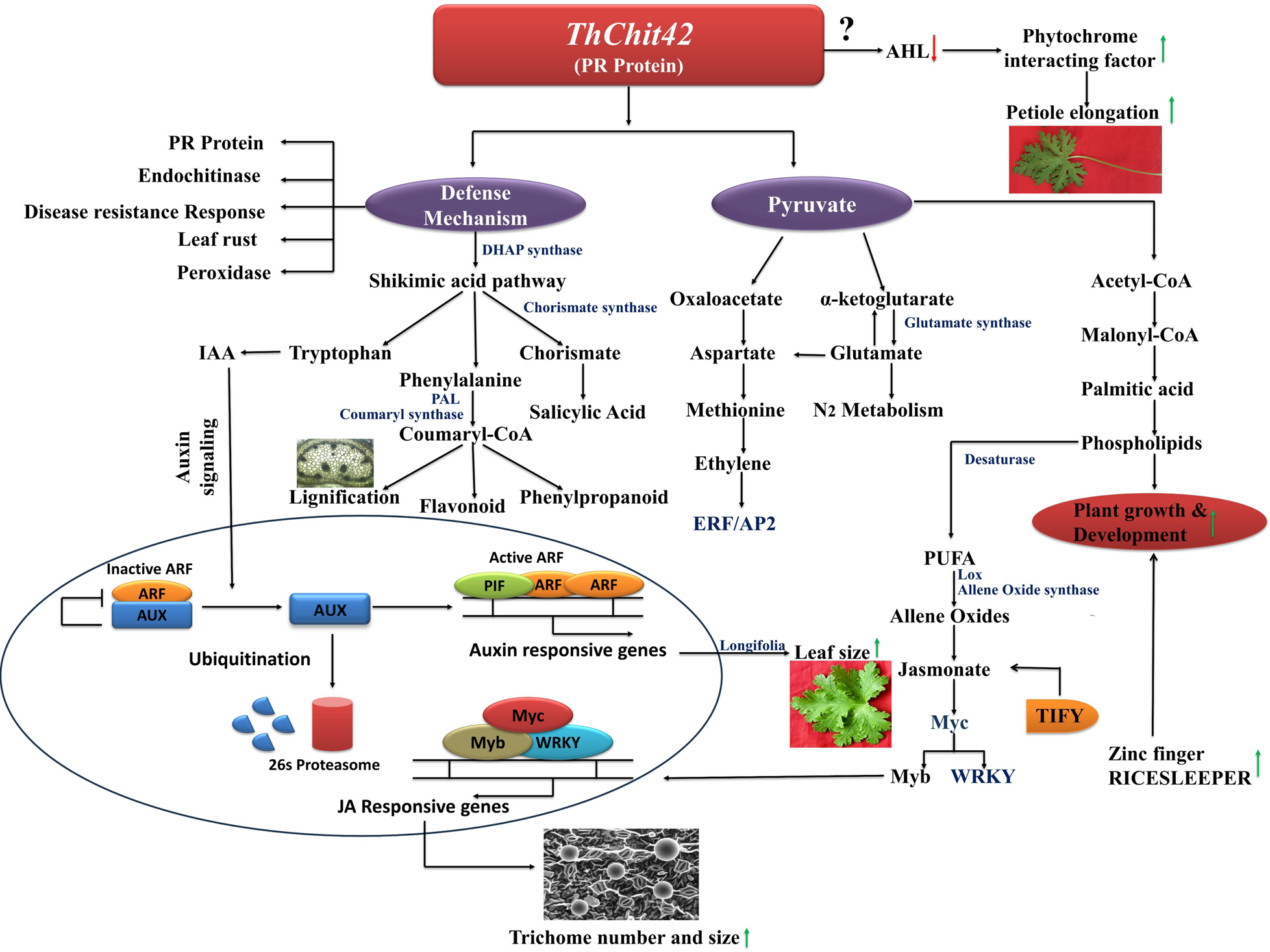
Hypothetical model proposed signal transduction linkage between growth and stress resistance pathway in sync with *Thchit42* gene from *Trichoderma harzianum*. Integration of 35S::*Thchit42* cassette in the *P. graveolens* plant genome led to overexpression of *Thchit42* (chitinase-PR protein), which resulted in a stress signal under the normal condition as the gene regulated by constitutive promoter. Plant cells upregulated defense mechanisms by rapidly producing *Pgchitinase, Pgendochitinase, PgDirigent (disease resistance response)*, and other stress resistance signalling pathways (JA) to overcome stress. Further, TIFY promotes downstream signalling (WRKY/MYC/TFs) for JA-responsive gene expression, which leads to enhanced trichomes. On the other hand, growth up-regulation is justified through glutamate synthase expression level. At the same time, auxin-regulated gene longifolia and AT-hook are responsible for a two-fold increase in leaf and petiole size. Transposons zinc finger BED domain RICESLEEPER intensive expression co-relates stress resistance with the growth of transgenic plants. Simultaneously, phenylpropanoid and flavonoid pathways have also been upregulated. See main text for additional details. Abbreviation: AHL, AT-hook motif nuclear-localized protein; PIF, Phytochrome Interacting Factor; PR-Pathogenesis Related protein; PAL, Phenylalanine Ammonia Lyase; ARF, Auxin Response Factor; AP2/ERF, Ethylene Response Factor.

## Discussion

Rose-scented Geranium is a pivotal contributor to the global GDP-contributing crops because of its vast application in the aromatherapy and perfumery industries. However, the growth and oil yield of the *P. graveolens* cultivar encounter substantial challenges during the rainy season in north India due to fungal stress, which hinder the global demand for GEO. Therefore, it is imperative for biotechnological approach to address fungal infection, ensuring the availability of propagules after the rainy season for further propagation (Lothe and Verma, 2023; Mazeed *et al.,* 2023; Varshney *et al.,* 2011). In this study, *P. graveolens* harboring 35S::*Thchit42* lines were developed for fungal resistance to maintain propagation rate and enhance yield. To develop fungal-resistant lines, mining of broad-spectrum PR protein was essential, resulting in *Thchit42* being screened based on transcript expression to keep the above promise (Carsolio *et al.,* 1994; Schlumbaum *et al.,* 1986) (Fig. S2). In addition, the comprehensive literature survey also suggests, *Thchit42* has considerable antifungal activity (Carsolio *et al.,* 1994). In our study, successful *Agrobacterium*-mediated genetic transformation was achieved with 70.0 ± 0.9% viability after acclimatization in a glass house (Fig. 1b) as per the previous report by Singh et al., (2021) Further, a 35S::*Thchit42* positive line L2 has shown significantly elevated relative expression of *Thchit42* among healthy propagating lines before and after biotic stress (Fig. 1d). Moreover, maximum transcript expression in L2 suggested that variation in the expression of the target gene among transformed lines was because of a combinatorial regulation system influences 35S promote (Warsi *et al.,* 2023).

In substantiating the molecular results, pathogenicity bioassays and antifungal activity have potentially contributed to disease resistance mechanism scoring. The transformed 35S:: *Thchit42* L2 and L4 plants showed 0 scoring, in contrast to the characteristic symptoms observed in the control plants when subjected to inoculation with *C. gloeosporioides* and *F. oxysporum*. This absence of disease symptoms in the 35S::*Thchit42* harboring lines undergoes enhanced resistance efficiency due to the over-expression of *Thchitinase*. The observed mitigation of disease symptoms in the transgenic line suggests augment chitinase level cannot be compensated for by other endogenous defense response genes in the native state (Fig. 3e-h). Remarkably, the recombinant *Thchit42* total protein showed promising antifungal activity, correlating with *in-vivo* and *in-vitro* leaf detached assay (Fig. 4a, c). A significant less leaf area necrosis depicts improved defence mechanism in transgenic plants compared to control (Lorito *et al.,* 1998). This efficacy may attribute to the direct hydrolysis of chitin and growing hyphae and the indirect involvement of chitin monomers in eliciting defense mechanisms. The findings presented herein align with the earlier reports (Chen *et al.,* 2014), corroborating and reinforcing the observed results.

Bolar et al., (2000) reported that expression of the *T. harzianum* endochitinase gene in Apple plants reduced growth and phenotype abnormalities. Meanwhile, the existing literature surveys on transgenic plants, i.e., cotton, tobacco, and potato with endochitinase expression, did not appear to have any adverse morphological anomalies (Lorito *et al.,* 1998). However, the present study revealed that the expression of 35S::*Thchit42* ameliorates phyllome growth along with increased trichome augmentation in transformed *P. graveolens* (Fig. 2b, d) (Fig. S8). Moreover, an increase in leaf and petiole size correlates positively with increased trichome numbers, showcasing clear association with an improved geranium essential oil content of ∼ 2.1fold, as shown in (Fig. 2e-g, i). In addition, histological analysis discloses an increased number of vascular bundles compared to control plants, which might explain the increment in petiole girth (Fig. 2c). Indeed, till the study, no known correlation was reported between *Thchit42* expression and vascular bundle number. Lucas et al., (2013) evaluated that tracheary elements (TEs), responsible for mineral and water transport, are characterized by secondary cell wall thickening. Moreover, the deposition of lignified secondary walls in TEs provides efficient water transport and avoids negative pressure during transpiration (Tu *et al.,* 2015). While a discrepancy in morphology and chemical composition content is not usual with endochitinase over-expression, the current findings add to the expanding literature on strategies for crop improvement. Interestingly, GEO has aromatic and medicinal properties; however, its production is compromised because of its unavailability after the rainy season. Hence, heterologous endochitinase gene transformation could be helpful to fulfil market demand sustainably.

Numerous studies support the successful genetic engineering to gain biotic and abiotic stress resistance in various crops, including tobacco, rice, carrot, geranium, and corn (Singh *et al.,* 2021; Patil and Widholm, 1997; Shukla *et al.,* 2016). However, no study reported genotype expression variability after integrating 35S::*Thchit42* in cultivar CIM-BIO171. Therefore, it is a privilege to unfold the considerable alteration in various gene expressions based on phenotype in transformed *P. graveolens* apart from fungal resistance (Fig. 2). The overall result hypothesized that *Thchit42* expression was constitutively present in the system as cassette 35S::*Thchit42* controlled by constitutive promoter 35S therefore, chitinase activity produces chitin oligosaccharides which act as potent elicitors for plant’s natural defense-related response directly or indirectly. Previously, defense-related changes were reported due to endochitinase expression, i.e., lignification, triggering ROS production (Fig. S9), and biosynthesis of jasmonic acid in other crops (Nojiri *et al.,* 1996) Still, the role of endochitinase in plant growth and essential oil content enhancement has yet to be reported. Hence, DEGs are a new addition to the list of genes known for participation in defense and growth mechanisms due to 35S::*Thchit42* integration. Interestingly, the indirect role of *Thchit42* in growth suggests that the target gene has a diverse role in plant-specialized mechanisms, as shown in Fig. 6; Fig. S8). Next, comprehensive literature strongly supports that all the up and down-regulated genes correlated through molecular mechanisms.

The observed up-regulation of *longifolia* and down-regulation of the AT-hook TFs postulated an alteration in leaf size and petiole elongation (Hwang *et al.,* 2017; Favero *et al.,* 2020). Additionally, for indirect defense response, the expression heatmap provides detailed information on the diversified role of *Thchit42* gene on coumaryl synthase, PAL, and caffeic acid-3-O-methyltransferase expression to provide lignification and activation of other disease-resistant or PR protein (Fig. S6, 7; Fig. 5c). Anisimova et al., (2021) and Li et al., (2017) had reported similar studies. Other findings suggest that PIF directly activated the auxin-responsive gene (IAA & YUC) to regulate *longifolia* for plant growth and development (Hwang *et al.,* 2017; Zhang *et al.,* 2022). Moreover, the over-expression of Lipoxygenase and allene oxide synthase, along with elevated TIFY for downstream signaling of MYC/WRKY/TFs, suggests a potential correlation with the presence of JA (He *et al.,* 2020). Chen et al., (2018) indicated that the up-regulation of JA possibly enhances trichome density for robust defense response. Although the defense response is constitutive, the growth of the transformed *P. graveolens* line remains uncompromised, as evidenced by the elevated glutamate synthase expression, which has a crucial role in nitrogen metabolism (Yu *et al.,* 2022). Further, the current study shows that glutamate also regulates ERF/AP2 levels to provide a strong resistance response (Fidler *et al.,* 2022). At the same time, a high transcript level of transposons Zinc Finger RICESLEEPER has unlocked the growth and resistance relationship (Knip *et al.,* 2012). Overall result is a model of *Thchit42* promising response towards development of fungal resistance transgenic crop in sync with plant growth and defence mechanism simultaneously. The current molecular mechanism findings could now be worthwhile to explore whether the model correlation has potential lead for other crops to achieve superior agronomic parameters for high economic benefits.

## Acknowledgments

The authors acknowledge the Council of Scientific and Industrial Research, India, for funding under the Aroma Mission Project (HCP0007), and the Council of Scientific and Industrial Research–Central Institute of Medicinal and Aromatic Plants (CSIR-CIMAP) for providing research facilities. The authors are deeply grateful to Director CSIR-CIMAP, Lucknow, for providing research platform to conduct all experiments. **Institutional communication number: CIMAP/PUB/2024/09**

## Competing Financial Interests

The authors declare, experiments were conducted in the absence of any commercial and financial relationships to avoid potential competing financial interests.

## Author contributions

K.K.: developed procedure to select *ThChit42* and screened, molecular cloning and expression of *ThChit42*, original manuscript preparation, acquisition of data, transcriptomic data evaluation and identification of potential gene to develop model co-relation. Z.I.W.: development of transgenic *Pelargonium graveolens* lines, execution of all experiments, critical manuscript editing, and acquisition of data. K.K., Z.I.W.: participated in all aspects of study, including molecular evaluation and phenotypic characterization of transgenic *P. graveolens* lines, hypothetical model correlation for genotype identification, localization of *ThChit42* through genome walking. A.S.: designing and supervision of pathogenicity bioassays and fungal activity. K.S.: designing and execution of pathogenicity bioassays for selection of fungal resistant transgenic lines, antifungal activity from total protein, execution of pathogen stress experiment for *ThChit42* selection. F.K., designing and supervision of *denovo* transcriptome *in-silico* study, critical manuscript editing. P.K.: execution of *denovo* transcriptome *in-silico* study. R.K.S.: genome walking supervision, critical manuscript editing. R.S.V.: essential oil chemical composition analysis. M.K.S.: execution of essential oil analysis. S.K.V.: statistical evaluation, Z.H.: review manuscript. G.P.: model designing and review manuscript. P.S.: scanning electron microscopic image. S.A.: review manuscript. L.U.R.: study concept and experimental design, study supervision, manuscript editing, critical review of experimental design, and intellectual input.

## Notes

### Competing Interest Statement

The authors have declared no competing interest.

## References

Afroz, S., Khatoon, K., Warsi, Z., Husain, Z., Verma, S. K. and Rahman, L. U. (2024) Molecular cloning and heterologous expression analysis of 1-Deoxy-D-Xylulose-5-Phosphate Synthase gene in *Centella asiatica* L. Gene 895, 148015.

Anisimova, O. K., Shchennikova, A. V., Kochieva, E. Z. and Filyushin, M. A. (2021) Pathogenesis-related genes of PR1, PR2, PR4, and PR5 families are involved in the response to Fusarium infection in garlic (*Allium sativum* L.). Int. J. Mol. Sci. 22, 6688.

Bergman, M. E., Chávez, Á., Ferrer, A. and Phillips, M. A. (2020) Distinct metabolic pathways drive monoterpenoid biosynthesis in a natural population of *Pelargonium graveolens*. J. Exp. Bot. 71, 258–271.

Bolar, J. P., Norelli, J. L., Wong, K. W., Hayes, C. K., Harman, G. E. and Aldwinckle, H. S. (2000) Expression of endochitinase from *Trichoderma harzianum* in transgenic apple increases resistance to apple scab and reduces vigor. Phytopathol. 90, 72–77.

Carsolio, C., Gutiérrez, A., Jiménez, B., Van Montagu, M. and Herrera-Estrella, A. (1994) Characterization of ech-42, a *Trichoderma harzianum* endochitinase gene expressed during mycoparasitism. Proc. Natl. Acad. Sci. U.S.A. 91, 10903–10907.

Chen, G., Klinkhamer, P. G., Escobar-Bravo, R. and Leiss, K. A. (2018) Type VI glandular trichome density and their derived volatiles are differently induced by jasmonic acid in developing and fully developed tomato leaves: implications for thrips resistance. Plant Sci. 276, 87–98.

Chen, P. J., Senthilkumar, R., Jane, W. N., He, Y., Tian, Z. and Yeh, K. W. (2014) Transplastomic *Nicotiana benthamiana* plants expressing multiple defence genes encoding protease inhibitors and chitinase display broad-spectrum resistance against insects, pathogens and abiotic stresses. Plant Biotechnol. J. 12, 503–515

Collinge, D.B., Kragh, K.M., Mikkelsen, J.D., Nielsen, K.K., Rasmussen, U. and Vad, K.(1993) Plant chitinases. Plant J. 3, 31–40.

Dana, M. D. L. M., Pintor-Toro, J. A. and Cubero, B. (2006) Transgenic tobacco plants overexpressing chitinases of fungal origin show enhanced resistance to biotic and abiotic stress agents. Plant Physiol. 142, 722–730.

Erb, M. and Kliebenstein, D. J. (2020) Plant secondary metabolites as defenses, regulators, and primary metabolites: the blurred functional trichotomy. Plant Physiol. 184, 39–52.

Fidler, J., Graska, J., Gietler, M., Nykiel, M., Prabucka, B., Rybarczyk-Płońska, A., Ewa, M., et al. (2022) PYR/PYL/RCAR receptors play a vital role in the abscisic-acid-dependent responses of plants to external or internal stimuli. Cells 11, 1352.

Favero, D. S., Kawamura, A., Shibata, M., Takebayashi, A., Jung, J. H., Suzuki, T., Jaeger, K. E. et al. (2020) AT-hook transcription factors restrict petiole growth by antagonizing PIFs. Curr. Biol. 30, 1454–1466.

Guarnaccia, V., Gilardi, G., Martino, I., Garibaldi, A. and Gullino, M. L. (2019) Species diversity in Colletotrichum causing anthracnose of aromatic and ornamental Lamiaceae in Italy. Agronomy 9, 613.

He, X., Kang, Y., Li, W., Liu, W., Xie, P., Liao, L., Luyao, H., et al. (2020) Genome-wide identification and functional analysis of the TIFY gene family in the response to multiple stresses in *Brassica napus* L. BMC Genom. 21, 1–13.

Horsch, R. B. (1985) A simple and general method for transferring genes into plants. Science 227, 1229–1231.

Howell, C. R. (2003) Mechanisms employed by Trichoderma species in the biological control of plant diseases: the history and evolution of current concepts. Plant Dis. 87, 4–10.

Hwang, G., Zhu, J. Y., Lee, Y. K., Kim, S., Nguyen, T. T., Kim, J. and Oh, E. (2017) PIF4 promotes expression of LNG1 and LNG2 to induce thermomorphogenic growth in Arabidopsis. Front. Plant Sci. 8, 1320.

Jiménez-Ortega, E., Kidibule, P. E., Fernández-Lobato, M. and Sanz-Aparicio, J. (2021) Structural inspection and protein motions modelling of a fungal glycoside hydrolase family 18 chitinase by crystallography depicts a dynamic enzymatic mechanism. Comput. Struct. Biotechnol. J. 19, 5466–5478.

Jogee, P. S., Ingle, A. P. and Rai, M. (2017) Isolation and identification of toxigenic fungi from infected peanuts and efficacy of silver nanoparticles against them. Food Control 71, 143–151.

Kalra, A., Parameswaran, T. N. and Ravindra, N. S. (1992) Fungicidal control of leaf-blight of geranium (*Pelargonium graveolens*). Indian J. Agric. Sci. 62, 844–847.

Knip, M., de Pater, S. and Hooykaas, P. J. (2012) The SLEEPER genes: a transposase-derived angiosperm-specific gene family. BMC Plant Biol. 12, 1–15.

Legrand, M., Kauffmann, S., Geoffroy, P. and Fritig, B. (1987) Biological function of pathogenesis-related proteins: four tobacco pathogenesis-related proteins are chitinases. Proc. Natl. Acad. Sci. U.S.A. 84, 6750–6754.

Li, N., Zhao, M., Liu, T., Dong, L., Cheng, Q., Wu, J., Le, W., et al. (2017) A novel soybean dirigent gene GmDIR22 contributes to promotion of lignan biosynthesis and enhances resistance to *Phytophthora sojae*. Front. Plant Sci. 8, 1185.

Lorito, M., Woo, S.L., Fernandez, I.G., Colucci, G., Harman, G.E., Pintor-Toro, J.A., Filippone, E., et al. (1998) Genes from mycoparasitic fungi as a source for improving plant resistance to fungal pathogens. Proc. Natl. Acad. Sci. U.S.A. 95, 7860–7865.

Lothe, N. B. and Verma, R. K. (2023) A study on geranium (*Pelargonium graveolens* L′ Herit ex Aiton) cultivars’ productivity and economics as intervening by diverse climatic conditions of the western peninsular region of India. Ind. Crops Prod. 200, 116882.

Lucas, W. J., Groover, A., Lichtenberger, R., Furuta, K., Yadav, S. R., Helariutta, Y. and Kachroo, P. (2013) The plant vascular system: evolution, development and functions. J. Integr. Plant Biol. 55, 294–388.

Luo, M., Xie, L., Chakraborty, S., Wang, A., Matny, O., Jugovich, M. and Ayliffe, M. (2021) A five-transgene cassette confers broad-spectrum resistance to a fungal rust pathogen in wheat. Nat. Biotechnol. 39, 561–566.

Mazeed, A., Maurya, P., Kumar, D., Prakash, O. and Suryavanshi, P. (2023) Enhancing productivity, quality, and economics of rose scented geranium (*Pelargonium graveolens* L.) through a novel integrated approach to phosphorus application. Ind. Crops Prod. 204, 117293.

Nojiri, H., Sugimori, M., Yamane, H., Nishimura, Y., Yamada, A., Shibuya, N. and Omori, T. (1996) Involvement of jasmonic acid in elicitor-induced phytoalexin production in suspension-cultured rice cells. Plant Physiol. 110, 387–392.

Patil, V. R. and Widholm, J. M. (1997) Possible correlation between increased vigour and chitinase activity expression in tobacco. J. Exp. Bot. 48, 1943–1950.

Rao, B. R. R. (2002) Biomass yield, essential oil yield and essential oil composition of rose-scented geranium (Pelargonium species) as influenced by row spacings and intercropping with cornmint (*Mentha arvensis* L.f. piperascens Malinv. ex Holmes). Ind. Crops Prod. 16, 133–144.

Schlumbaum, A., Mauch, F., Vögeli, U. and Boller, T. (1986) Plant chitinases are potent inhibitors of fungal growth. Nature 324, 365–367.

Shukla, A. K., Upadhyay, S. K., Mishra, M., Saurabh, S., Singh, R., Singh, H., Thakur, N., et al. (2016) Expression of an insecticidal fern protein in cotton protects against whitefly. Nat. Biotechnol. 34, 1046–1051.

Singh, P., Pandey, S. S., Dubey, B. K., Raj, R., Barnawal, D., Chandran, A. and Rahman, L. U. (2021) Salt and drought stress tolerance with increased biomass in transgenic *Pelargonium graveolens* through heterologous expression of ACC deaminase gene from *Achromobacter xylosoxidans*. Plant Cell, Tissue Organ Cult. 147, 297–311.

Srivastava, P., Garg, A., Misra, R. C., Chanotiya, C. S. and Ghosh, S. (2021) UGT86C11 is a novel plant UDP-glycosyltransferase involved in labdane diterpene biosynthesis. J. Biol. Chem. 297.

Stukenbrock, E. and Gurr, S. (2023) Address the growing urgency of fungal disease in crops. Nature 617, 31–34.

Tu, B., Hu, L., Chen, W., Li, T., Hu, B., Zheng, L., and Li, S. (2015) Disruption of OsEXO70A1 causes irregular vascular bundles and perturbs mineral nutrient assimilation in rice. Sci. Rep. 5, 18609.

Upadhyay, R., Lothe, N. B., Bawitlung, L., Singh, S., Singh, M. K., Kumar, P., Verma, K. R., et al. (2022) Secondary metabolic profile of rose-scented geranium: A tool for characterization, distinction and quality control of Indian genotypes. Ind. Crops Prod. 187.

Varshney, R. K., Bansal, K. C., Aggarwal, P. K., Datta, S. K. and Craufurd, P. Q. (2011) Agricultural biotechnology for crop improvement in a variable climate: hope or hype? Trends Plant Sci. 16, 363–371.

Warsi, Z. I., Khatoon, K., Singh, P. and Rahman, L. U. (2023) Enhancing drought resistance in *Pogostemon cablin* (Blanco) Benth. through overexpression of ACC deaminase gene using thin cell layer regeneration system. Front. Plant Sci. 14.

Yu, B., Liu, N., Tang, S., Qin, T. and Huang, J. (2022) Roles of Glutamate Receptor-Like Channels (GLRs) in Plant Growth and Response to Environmental Stimuli. Plants 11, 3450.

Zhang, Y., Yu, J., Xu, X., Wang, R., Liu, Y., Huang, S., Hairong, W., et al. (2022) Molecular mechanisms of diverse auxin responses during plant growth and development. Int. J. Mol. Sci. 23, 12495.

