## Supplementary file for "Bridging fungal resistance and plant growth through constitutive overexpression of *Thchit42* gene in *Pelargonium graveolens*"

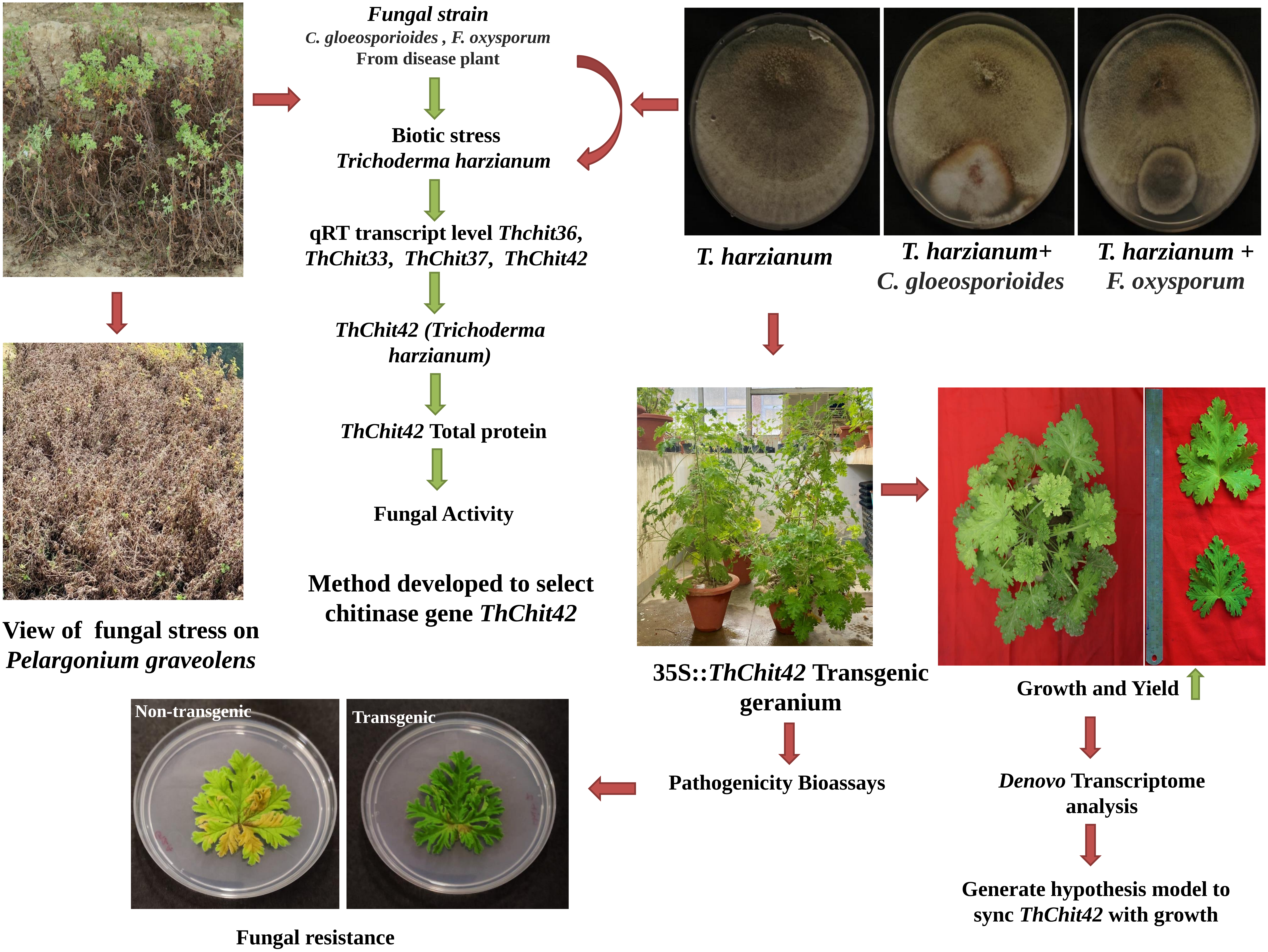

Fungal strain
C. gloeosporioides , F. oxysporum
From disease plant
Biotic stress
Trichoderma harzianum
qRT transcript level Thchit36, ThChit33, ThChit37, ThChit42
ThChit42 (Trichoderma harzianum)
ThChit42 Total protein
Fungal Activity
Method developed to select chitinase gene ThChit42
T. harzianum
T. harzianum+ C. gloeosporioides
T. harzianum + F. oxysporum
View of fungal stress on Pelargonium graveolens
35S::ThChit42 Transgenic geranium
Growth and Yield
Pathogenicity Bioassays
Denovo Transcriptome analysis
Generate hypothesis model to sync ThChit42 with growth
Non-transgenic
Transgenic
Fungal resistance

**Supplementary Figure 1**

**Outline of the work**

Simplified representation of harmonizing fungal resistance with growth

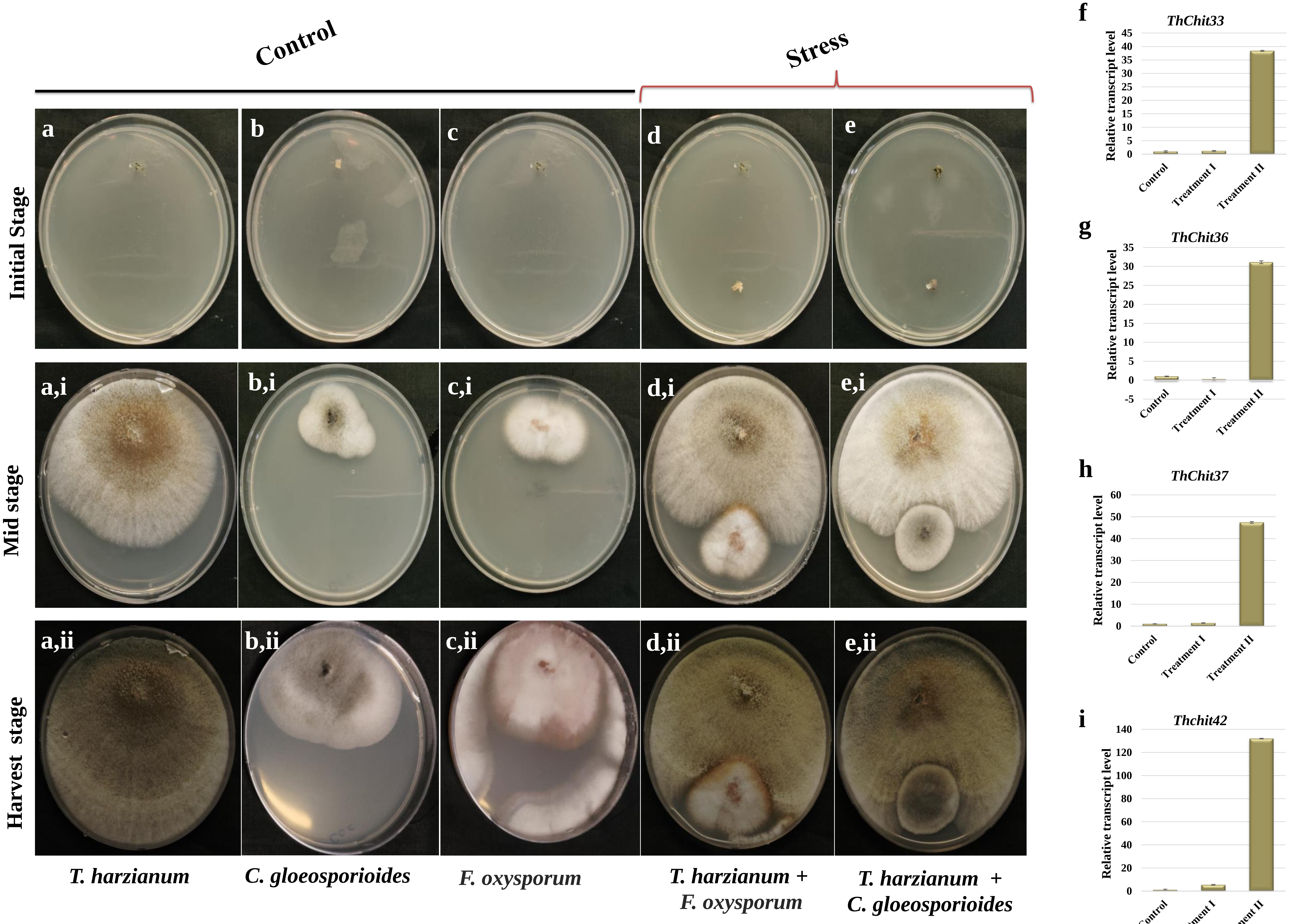

Control
Stress
Initial Stage
Mid stage
Harvest stage
a
b
c
d
e
a,i
b,i
c,i
d,i
e,i
a,ii
b,ii
c,ii
d,ii
e,ii
T. harzianum
C. gloeosporioides
F. oxysporum
T. harzianum + F. oxysporum
T. harzianum + C. gloeosporioides
f
ThChit33
g
ThChit36
h
ThChit37
i
Thchit42
Relative transcript level
Control
Treatment I
Treatment II

**Supplementary Figure 2**

Procedure to select *Thchitinase* gene from *Trichoderma harzianum* based on relative transcript expression under *C. gloeosporioides* and *F. oxysporum* stress. **(a, i, ii)** *Trichoderma harzianum* control plate at different stages of growth. **(b, i, ii)** *C. gloeosporioides* control culture growth. **(c, i, ii)** *F. oxysporum* control at different growth intervals. **(d, i, ii)** Aggressive fungal disc of *F. oxysporum* inoculated with *T. harzianum* for mRNA relative expression evaluation of diverse chitinase. **(e, i, ii)** *C. gloeosporioides* inoculated with *T. harzianum* to analyse various chitinase relative transcript expressions. **(f-i)** mRNA expression level of *Thchit33*, *Thchit36, Thchit37*, and *Thchit42* from *Trichoderma harzianum*. Experiments were performed in 3 technical repeats; error bars, mean ± s.d (n=3)

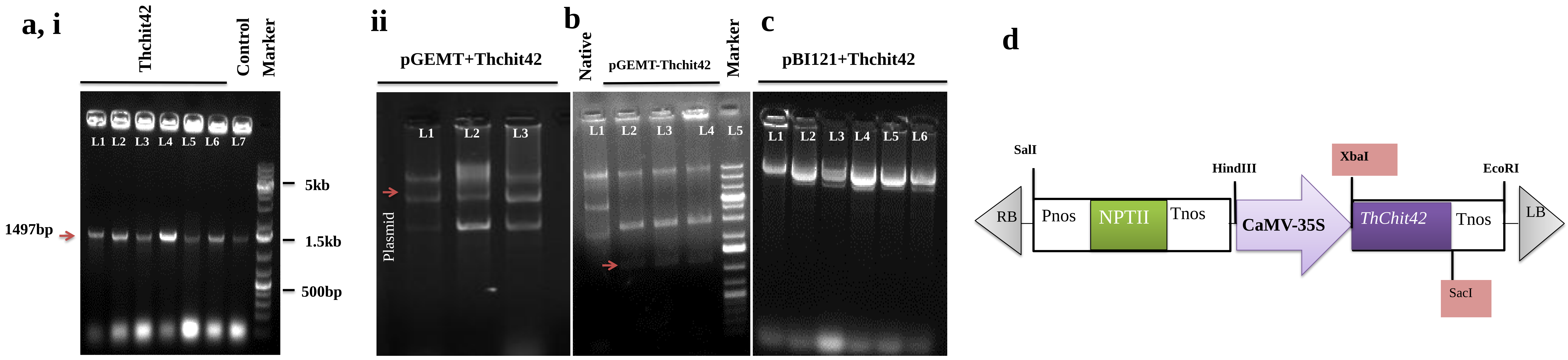

a, i
Thchit42
Control
Marker
L1 L2 L3 L4 L5 L6 L7
1497bp
5kb
1.5kb
500bp
ii
pGEMT+Thchit42
L1 L2 L3
Plasmid
b
Native
pGEMT-Thchit42
Marker
L1 L2 L3 L4 L5
c
pBI121+Thchit42
L1 L2 L3 L4 L5 L6
d
SalI
HindIII
XbaI
EcoRI
RB
Pnos
NPTII
Tnos
CaMV-35S
ThChit42
Tnos
LB
SacI

**Supplementary Figure 3**

Molecular cloning of endochitinase *Thchit42* from *T. harzianum*. **(a, i, ii)** Positive clones and plasmid of *Thchit42* in pGEMT. **(b)** Clones were confirmed through restriction digestion with SacI and XbaI sticky end cutters. **(c)** The purified product was cloned in the pBI121 vector under the constitutive promoter CaMV35S.lane, 1kb + ladder. **(d)** Schematic diagram of T-DNA cassette having *nptII* (Kanamycin resistant) and 35S::*Thchit42* gene.

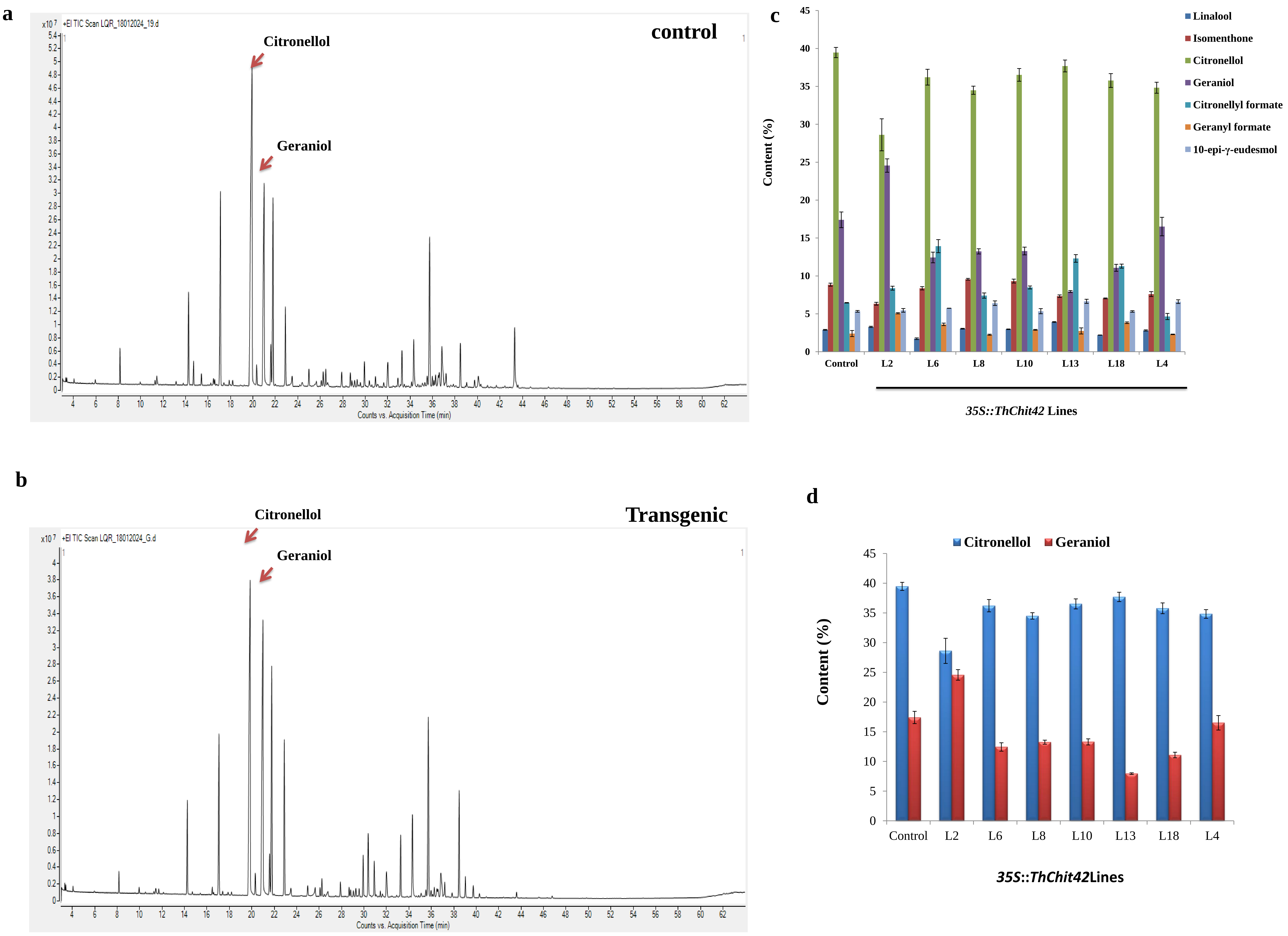
a
x10 7 +EI TIC Scan LQR_18012024_19.d
control
Citronellol
Geraniol
Counts vs. Acquisition Time (min)
b
x10 7 +EI TIC Scan LQR_18012024_G.d
Transgenic
Citronellol
Geraniol
Counts vs. Acquisition Time (min)
c
Linalool
Isomenthone
Citronellol
Geraniol
Citronellyl formate
Geranyl formate
10-epi-γ-eudesmol
Content (%)
Control
L2
L6
L8
L10
L13
L18
L4
35S::ThChit42 Lines
d
Citronellol
Geraniol
Content (%)
Control
L2
L6
L8
L10
L13
L18
L4
35S::ThChit42Lines

**Supplementary Figure 4**

Representation of *Pelargonium graveolens* essential oil composition. **(a, b)** GC-MS chromatograms of control (CIM-B171) and transgenic line L2 essential oil. **(c)** Chemical composition content of transgenic lines and control. (d) Representation of citronellol and geraniol content. Experiments were performed in 3 technical repeats; error bars, mean ± s.d (n=3)

a

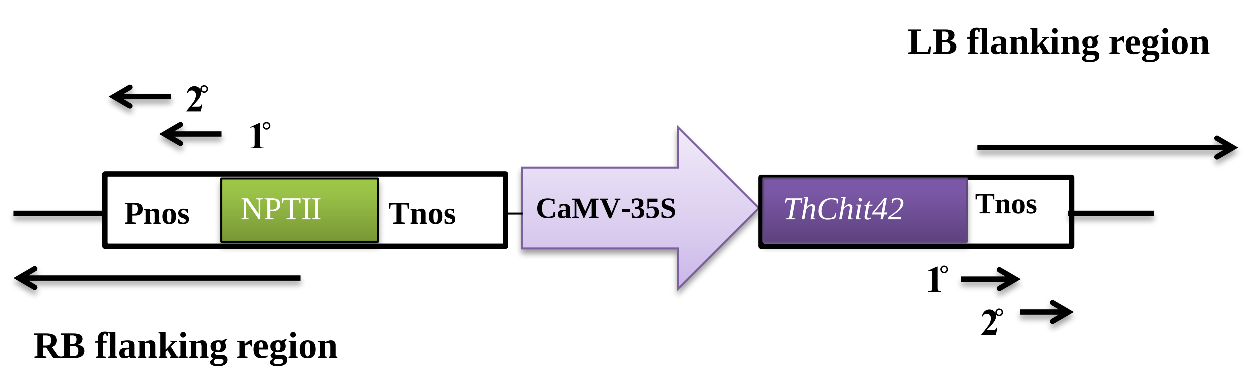
LB flanking region
2°
1°
Pnos
NPTII
Tnos
CaMV-35S
ThChit42
Tnos
1°
2°
RB flanking region

**Supplementary Figure 5**

**(a)** Schematic diagram of T-DNA cassette for genome walking. Short arrows indicate primary and nested primers; long arrows indicate RB and LB flanking regions.

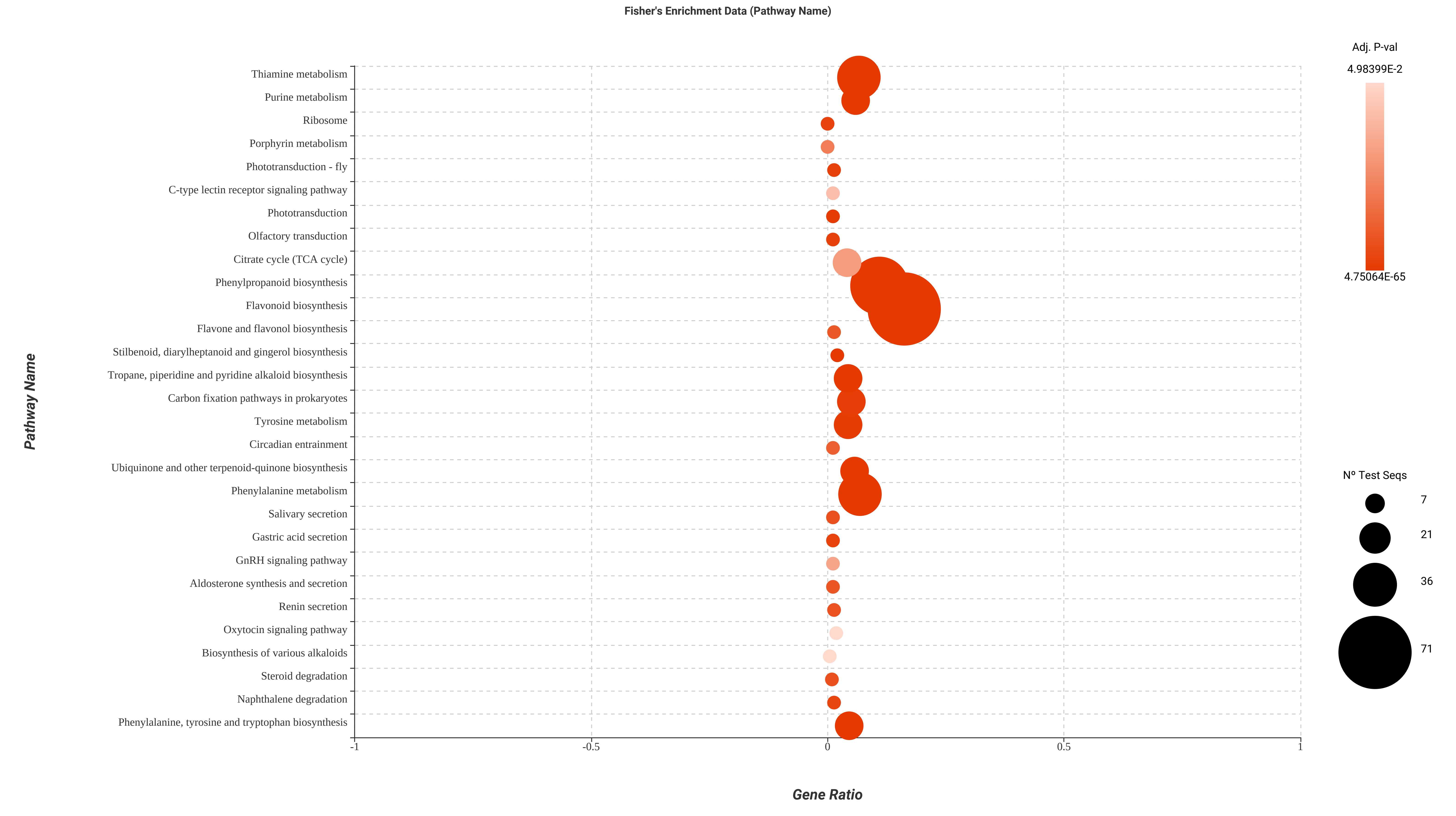
Fisher's Enrichment Data (Pathway Name)
Pathway Name
Thiamine metabolism
Purine metabolism
Ribosome
Porphyrin metabolism
Phototransduction - fly
C-type lectin receptor signaling pathway
Phototransduction
Olfactory transduction
Citrate cycle (TCA cycle)
Phenylpropanoid biosynthesis
Flavonoid biosynthesis
Flavone and flavonol biosynthesis
Stilbenoid, diarylheptanoid and gingerol biosynthesis
Tropane, piperidine and pyridine alkaloid biosynthesis
Carbon fixation pathways in prokaryotes
Tyrosine metabolism
Circadian entrainment
Ubiquinone and other terpenoid-quinone biosynthesis
Phenylalanine metabolism
Salivary secretion
Gastric acid secretion
GnRH signaling pathway
Aldosterone synthesis and secretion
Renin secretion
Oxytocin signaling pathway
Biosynthesis of various alkaloids
Steroid degradation
Naphthalene degradation
Phenylalanine, tyrosine and tryptophan biosynthesis
-1
-0.5
0
0.5
1
Gene Ratio
Adj. P-val
4.98399E-2
4.75064E-65
Nº Test Seqs
7
21
36
71

**Supplementary Figure 6**

Significantly enriched pathways based on FDR < 0.05. Refer Figure 5 and 6 to correlate pathways.

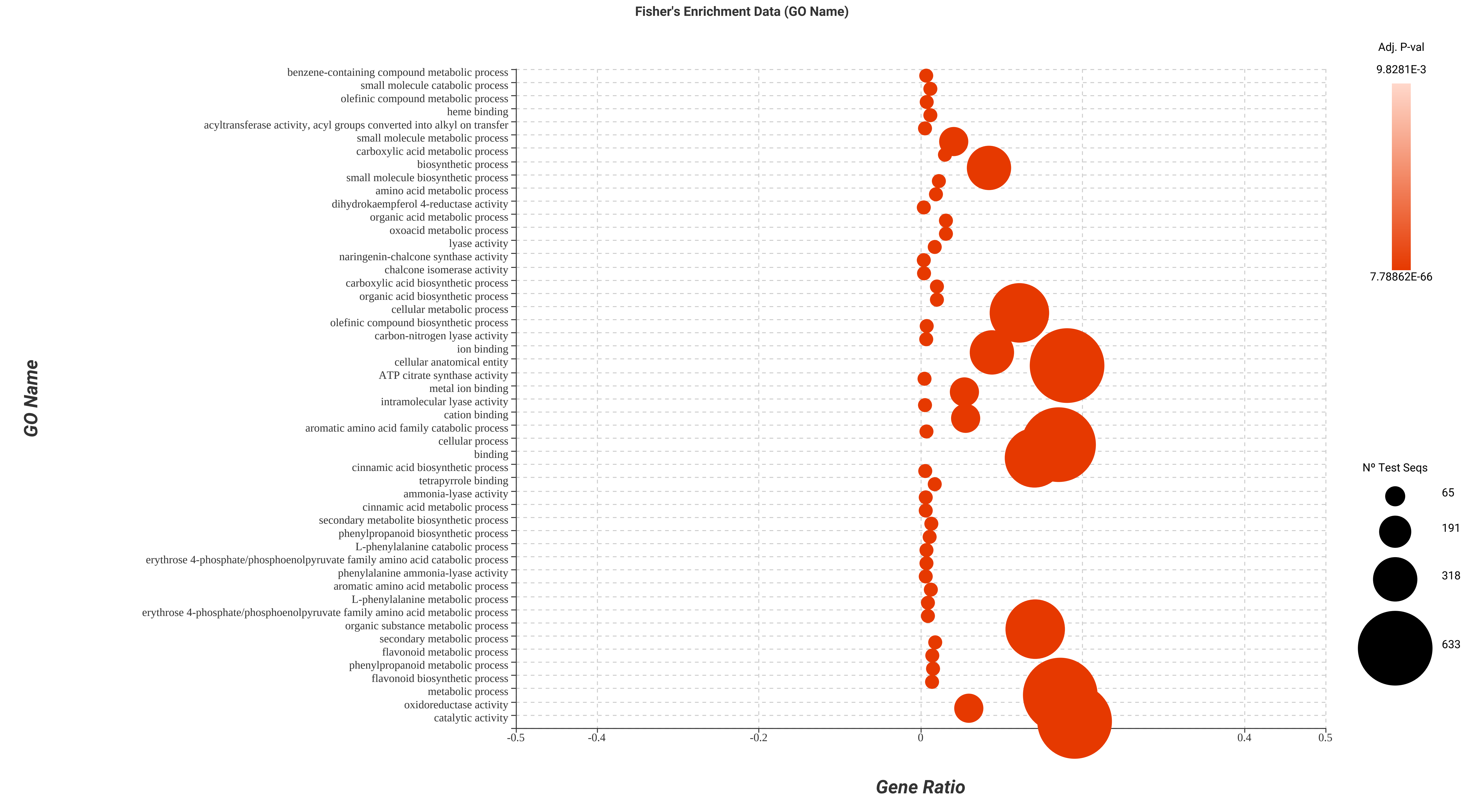
Fisher's Enrichment Data (GO Name)
GO Name
benzene-containing compound metabolic process
small molecule catabolic process
olefinic compound metabolic process
heme binding
acyltransferase activity, acyl groups converted into alkyl on transfer
small molecule metabolic process
carboxylic acid metabolic process
biosynthetic process
small molecule biosynthetic process
amino acid metabolic process
dihydrokaempferol 4-reductase activity
organic acid metabolic process
oxoacid metabolic process
lyase activity
naringenin-chalcone synthase activity
chalcone isomerase activity
carboxylic acid biosynthetic process
organic acid biosynthetic process
cellular metabolic process
olefinic compound biosynthetic process
carbon-nitrogen lyase activity
ion binding
cellular anatomical entity
ATP citrate synthase activity
metal ion binding
intramolecular lyase activity
cation binding
aromatic amino acid family catabolic process
cellular process
binding
cinnamic acid biosynthetic process
tetrapyrrole binding
ammonia-lyase activity
cinnamic acid metabolic process
secondary metabolite biosynthetic process
phenylpropanoid biosynthetic process
L-phenylalanine catabolic process
erythrose 4-phosphate/phosphoenolpyruvate family amino acid catabolic process
phenylalanine ammonia-lyase activity
aromatic amino acid metabolic process
L-phenylalanine metabolic process
erythrose 4-phosphate/phosphoenolpyruvate family amino acid metabolic process
organic substance metabolic process
secondary metabolic process
flavonoid metabolic process
phenylpropanoid metabolic process
flavonoid biosynthetic process
metabolic process
oxidoreductase activity
catalytic activity
-0.5
-0.4
-0.2
0
0.4
0.5
Gene Ratio
Adj. P-val
9.8281E-3
7.78862E-66
Nº Test Seqs
65
191
318
633

**Supplementary Figure 7**

Significantly enriched gene ontology terms based on FDR < 0.05. Refer Figure 5 and 6 to correlate pathways.

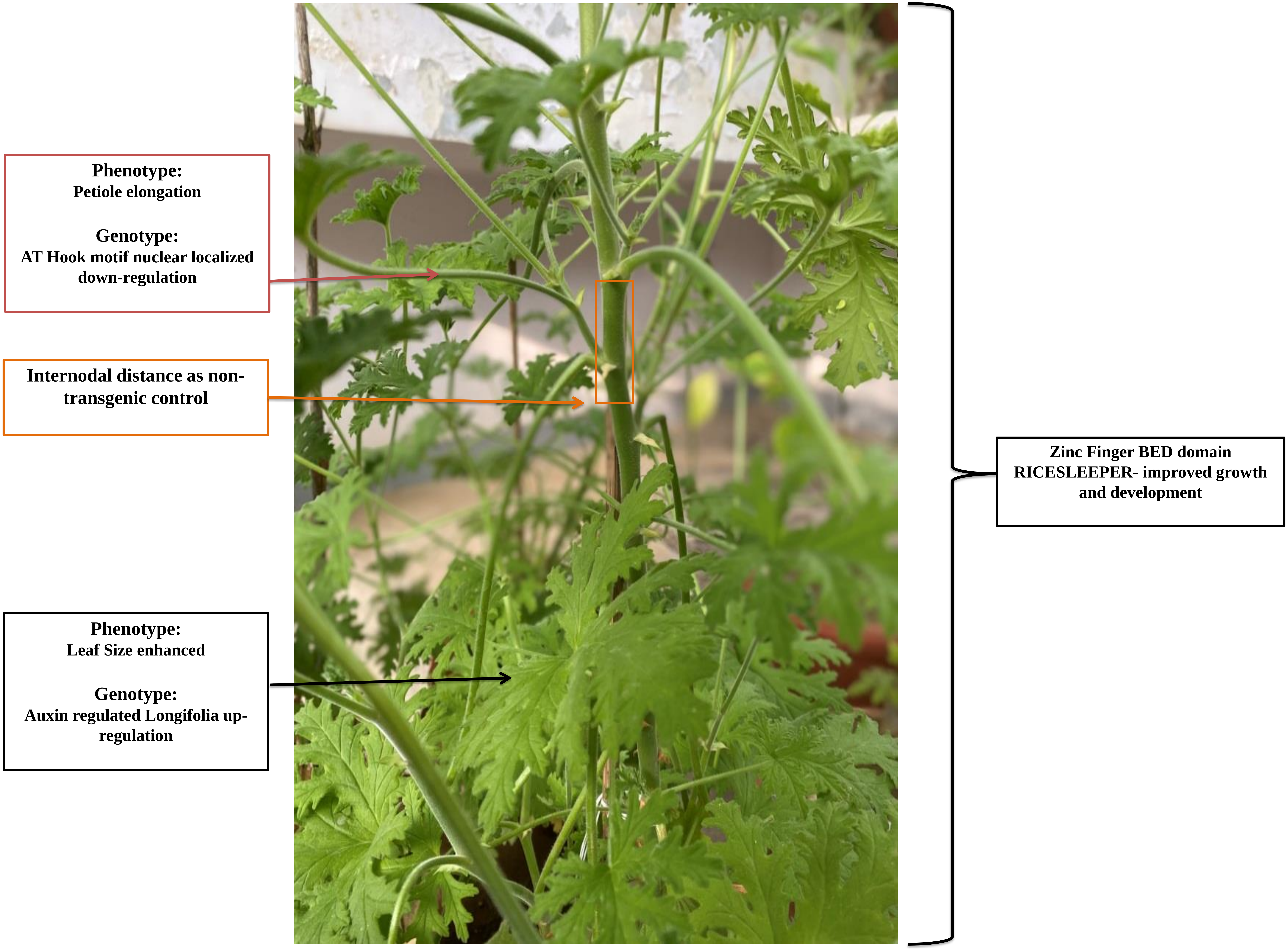
Phenotype:
Petiole elongation
Genotype:
AT Hook motif nuclear localized down-regulation
Internodal distance as non-transgenic control
Phenotype:
Leaf Size enhanced
Genotype:
Auxin regulated Longifolia up-regulation
Zinc Finger BED domain RICESLEEPER- improved growth and development

**Supplementary Figure 8**

Co-ordination of forward and reverse genetics for growth improvement

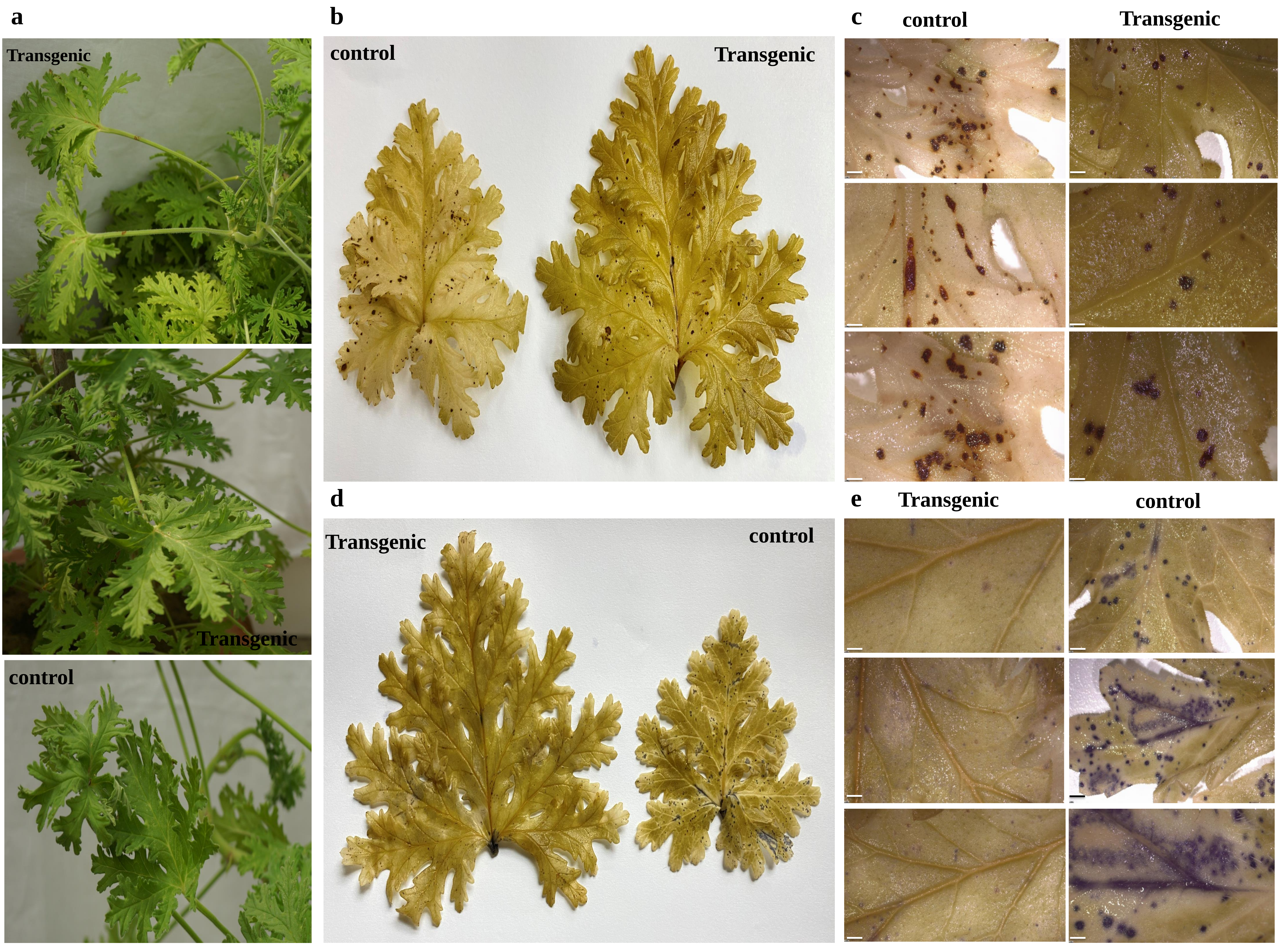
a
Transgenic
Transgenic
control
b
control
Transgenic
c
control
Transgenic
d
Transgenic
control
e
Transgenic
control

**Supplementary Figure 9**

Estimation of ROS ($H_2O_2$ and $O_2^-$) generation under *C. gloeosporioides* and *F. oxysporum* potent fungal stress. **(a)** Fungal inoculation in the leaf of transgenic and control plants. **(b)** 3, 3'- Diaminobenzidine (DAB) spots on transgenic and control leaves indicate less accumulation of $H_2O_2$ (brown spots) in transgenic leaves under biotic stress. (c) Stereomicroscopic visualization of DAB spots. **(d)** Nitro blue tetrazolium (NBT) stained (blue spots) evaluates amplified $O_2^-$ in control leaf contrast to transgenic leaf under fungal stress. **(e)** Microscopic view of NBT spots, examined under a stereomicroscope. Experiments were performed thrice in 3 technical repeats.

**Supplementary Table 1** *Denovo* transcriptome data analysis

| | CL | WT |
|---|---|---|
| **Raw reads** | 32045514 | 26806888 |
| **Assembled transcripts** | 234557 | 237989 |
| **N50** | 726.62 | 850.78 |
| **Remapping % (Bowtie2)** | 86.77 | 79.61 |
| **BUSCO %** | 76.71 | 97.88 |
| **CD-HIT-EST (unigenes)** | 263402 | |
| **Blastx** | 133809 | |
| **GO mapped** | 49017 | |
| **KEGGs** | 343 | |
| **DEGs** | 3328 (2158 upregulated; 1170 downregulated) | |

CL- 35S::*Thchit42* transgenic line L2; WT- non-transgenic control

**Supplementary Table 2** List of primers used in PCR and qRT-PCR

| Gene | Primer sequence (5´- 3´) | Accession number |
|---|---|---|
| *nptII* | F-AAGATGGATTGCACGCAGGT<br>R-TCAGAAGAACTCGTCAAGAAGGC | [19] (AF485783) |
| *Thchit42* | F-ATGTTGAGCTTCCTGGGCAA<br>R-TTAGTTCAGGCCGTTCTTAATG | S78423 |
| qRT *Thchit42* | F-TCACGACGCCAACCTGTTTA<br>R-TAGATGGGCATGCCGAGAAC | S78423 |
| qRT *Pgendochit* | F- GGAAGCCAAACTCCATGTCC<br>R- TCAAGAGGACTAGGTGCAGC | - |
| qRT *Pgchitinase* | F- TCTGTGGCACTGGTGAAGAA<br>R- TTGAAGAAATCCGGCGTCAC | - |
| qRT *PgDirigent* | F- AAATGCTCACAACGCCACAT<br>R- ATTGGGTCGTCGAAAACAGC | - |
| qRT *PgWRKY* | F- GGCTCCCAGTTGTCCAGTTA<br>R- CTCGTCTCCCAAGCATCTCT | - |
| qRT *PgFMO* | F- GCATGAAGTGATCCGAGTGG<br>R- ATCCGCAAGTTTTGGCTCAG | - |
| qRT *PgAOS* | F- AAACACGCAGCTCTCAAAGG<br>R- CAGAGTAGTCGGCTTTCCCA | - |
| qRT *PgAHL* | F- CAATTCGCCAACCTTGCTCT<br>R- CCGCCGGTTCTACTCCTTAT | - |
| qRT *PgLongifolia* | F- AGATAGATGGAAGGCGAGGC<br>R-ACCGCTTCTCAATCTCACCA | - |
| qRT *PgARF* | F- AGGTTGGCTGGGATGAATCA<br>R- CCTTGGTCGCTTGGATCTGA | - |
| qRT *PgZF* | F- TGATGTATCCTTCGAACTCTGC<br>R- CCACCAACCACTTGAGAAGG | - |
| qRT *PgG6PD* | F- TAAAGGCCGGGAAAGCACTA<br>R- GGGTTGCAGGCGAATTACAA | - |
| qRT *PgAct* | F- AATCTTCTTGCCGCTCATGC<br>R- AGATTGTGGACAGCTGCTCT | - |
| qRT *PgAP2/ERF* | F- TACTCCTTTCCCTGCATCCC<br>R- ACCAAGTCTCTGAACACCGA | - |
| qRT *PgGS* | F- GTTGGGTCATAACGGGGAGA<br>R- CATTCTTCGAGAGGCCAAGC | - |
| qRT *PgTIFY* | F- GCCGGAAAGCATGTTAGGTC<br>R- AGGGGCCGGCATAATATCTG | - |
| qRT *ThAct* | F- ACTGGGACGACATGGAGAAG<br>R- CTGGGTCATCTTCTCACGGT | AM231150 |
| qRT *ThTub* | F- TGGACGAGATGGAGTTCACC<br>R- CTCCTCCTCGTACTCCTCCT | MK895942 |
| qRT *ThChit36* | F- GGAAAACGACATGGCTCCTG<br>R- AGAAGCACCTCCGATTGACA | AY028421 |
| qRT *ThChit37* | F-GTTCACCCCGGATTTGGATG<br>R-ATCCATGCCATCTTCCCACA | AF525753 |
| qRT *ThChit33* | F- ACCAGCACCCAGAACAACTA<br>R-TTGAGAGCCAGAGACGTAGC | X80006 |

F - Forward, R – Reverse, GS – Glutamate Synthase, ZF – Zinc Finger BED domain RICESLEEPER, AHL – AT Hook motif nuclear Localization, G6PD – Glucose-6-Phosphate Dehydrogenase, ERF – Ethylene Response Factor, FMO – Flavin Monooxygenase, AOS – Allene Oxide Synthase

**Pg – Pelargonium graveolens*

**Th – Trichoderma harzianum*

**Supplementary Table 3** List of primers used in Genome walking

| Gene | Primer sequence (5´- 3´) |
|---|---|
| Pnos | R1-GAACGCGCAATAATGGTTTCTGACGTATG<br>R2-TGAGTGGCTCCTTCAACGTTGCGGTTCTG |
| Tnos | R1- TAATCATCGCAAGACCGGCAACAGGATTC<br>R2- TGCTGCAAGGCGATTAAGTTGGGTAACGC |

Pnos- Nos promoter; Tnos- Nos terminator; R1- Primary reverse primer; R2- Nested reverse primer

**Supplementary Table 4** Pathogenicity bioassays of *Colletotrichum gloeosporioides* seven days post inoculation on *P. graveolens* transgenic and non-transgenic leaves

| **S. No** | **Transgenic lines** | **Pathogenicity test inference** |
|---|---|---|
| 1 | W | H |
| 2 | Line 1 | H |
| 3 | Line 2 | O |
| 4 | Line 3 | M |
| 5 | Line 4 | O |
| 6 | Line 5 | L |
| 7 | Line 6 | O |
| 8 | Line 7 | H |
| 9 | Line 8 | O |
| 10 | Line 9 | M |
| 11 | Line 10 | H |
| 12 | Line 11 | L |
| 13 | Line 12 | O |
| 14 | Line 13 | O |
| 15 | Line 14 | O |
| 14 | Line 15 | O |

W- Non-transgenic control; **Line**- Transgenic lines; **O**- healthy plant; **L-** presence of low virulent lesion; M -moderate lesion; **H-**high presence and spread of necrotic spot

**Supplementary Note 1**

**Localization of T-DNA cassette site**

**RB flanking sequence of T-DNA cassette (35S::*ThChit42*)**

AAGGCGGAAGGGCCCCTGGGGCGCCGGGAAACCCCGCCAGTTACAAATTATACCAAAACCCCCCA
ACAACGGCGACGTTCCAAAACTAATAATGTTTTTTTGTCCCCAACAATGGTTGGGTTTTTAAAAAG
TGGGGTTTTTTCTCCCTCAAAGTGCCGTTCCCTTCCTCCCCAAAAAAAATATTATCAATTTTGGGCC
CGGGTGGGTCGCCCCCCCGGGTTTAAACCGGAGGGGGAAAAACAACATTCTTTCTGGGCGGAAAT
TTTAAGGAACCTTTTTATCCCCCCGCGATGTCACGCAACACGAGCTTCTTTAAACC

ATATAGTAGTTGTTGGATACTCTTTTTTTTTTTTTTTATCGCCTACACACCCCGCGCCTTTTTATATAT
GTGGGGTGAGAATATTCGGGAGAGCCCTCATTTTGGGGGTTTTTGGGGGCCCCTTTTTTCCCCCCCC
AAAAAAAAAAAAAACCCCCCCCCCCACAAAAAAAAAAAACTTTTCTTTGTGTGTGCTTTTGGCGAA
AAAACCAGAGGGGGGTTTTTTTATTTGGGAATTTTCCCCCGGTTTTTCGGTTGGGCTTCCCCCCAGG
GGGGAATTTTTGGGGGGGGGGGGCTTCTTTTTGGCTTCTTTTTTCCCCCTCGGCCCCCCTTTAAAA
AAAGAGAGAGAGGGTTTTTTTTGGGGGGGGTTTTTTTCCCCCCAAAAAAAAAAAAAAGGGCTTTTTT
TAAAAAAAAAAAAAAAGGAAATTTTTTTTCCGCCCCCTTTTTTTTTGGGGGGGGGGTTTTTTTTTTT
GTTTTAAAAAAGGGGCCCCCCTTTTTTTTTTGTTTTTTTTGGGGGGGAGATTCTTTTTAGGGGGCGCC
ACTAGGAAGATTTTTCTTTTTTCGGGGAACTTGAAAATGGGTAAAATTAATATTCTAAAGGGGGAT
CTTTTTTTTTTAAGGTTGGAATTTAAAAGGAAAAAAATGTTTTTTTCCGTTTTTGGGGGGCGGCCTG
AATGGGGGGAAAAAAAAATTATTCCTTCCCCAAAAACCCTTAAGGAACAATTTTTTAAAAAAAAA
AAAAAACGCCCCCCCGGACCAACTCCTAAATTGGGGGGGGGGACCAGGATTCCCCCCCCACCCCT
TCCTTCTCGGGGGAAAAAAACCCCACAAAAACTCCTTCCTCGGGGGGGGGGGGGGGCTCCCAAAA
ACCCCCCCCCCCCTCCATCCAACCATTATTGTTTAAACCGGAGGGCGGGAACCACCATTCTTTCGG
GCGGAAGAAAAAGGGTCCTCGTTCTCCCCCCGCGATCGACGCGACACACGCTCTTTTGAAAT

Black line – TATA and CAAT sequence; Red Line- multiples of T and A; Green line multiples of C

An interpretation of heterologous gene insertion sites within the plant genome is prerequisite for a possible explanation of transgene integration position. The erroneous insertion of a T-DNA cassette might have uncertain effects, i.e., silencing or overexpression of endogenous or non-endogenous gene (Cullen *et al.*, 2011). Notably, *Agrobacterium*-mediated T-DNA transfer occurs explicitly at the RB and ends at the LB (Tiwari *et al.*, 2022). Therefore, integration site investigation of construct in the genomic DNA of transformed L2, the reverse primer of Pnos and Tnos was designed for genome walking. Likewise, Li et al. (2015) demonstrated that primary and nested PCR (partial-overlapping) based genome walking correlate with particular amplified products. Consequently, the results evaluated LB and RB flanking sequences from genome walking libraries of DraI and PvuII which do not hamper the expression of other genes as cassettes have integrated at non-coding regions (Supplementary note 1). Rajpriya et al. (2021) reported a similar T-DNA localization study. As per the literature study, both the sequence has TATA and CAAT regions for transcription. Additionally, it represents multiples of all four nucleotides which might denote intron

(Tsuritani et al., 2007; Yella et al., 2018). Furthermore, the sequence validated that T-DNA insertion at the intron has not caused a position effect on other gene expressions.

**Supplementary references**

Cullen, D., Harwood, W., Smedley, M., Davies, H. and Taylor, M. (2011) Comparison of DNA walking methods for isolation of transgene-flanking regions in GM potato. *Mol. Biotechnol.* **49**, 19-31.

Li, H., Ding, D., Cao, Y., Yu, B., Guo, L. and Liu X. (2015) partially overlapping primer-based PCR for genome walking. PLoS One 10, e0120139.

Rajapriya, V., Kannan, P., Sridevi, G. and Veluthambi, K. (2021) A rare transgenic event of rice with Agrobacterium binary vector backbone integration at the right T-DNA border junction. *J. Plant Biochem. Biotechnol.* 1-8.

Tiwari, M., Mishra, A. K. and Chakrabarty, D. (2022) Agrobacterium-mediated gene transfer: recent advancements and layered immunity in plants. Planta 256, 37.

Tsuritani, K., Irie, T., Yamashita, R., Sakakibara, Y., Wakaguri, H., Kanai, A., Mizushima-Sugano, J. *et al*. (2007) Distinct class of putative "non-conserved" promoters in humans: comparative studies of alternative promoters of human and mouse genes. *Genome Res.* **17**, 1005-1014.

Yella, V. R., Kumar, A. and Bansal, M. (2018) Identification of putative promoters in 48 eukaryotic genomes on the basis of DNA free energy. *Sci. Rep.* **8**, 4520.
